# A Novel Biosensor Reveals Dynamic Changes of C-di-GMP in Differentiating Cells with Ultra-High Temporal Resolution

**DOI:** 10.1101/2022.10.18.512705

**Authors:** Andreas Kaczmarczyk, Simon van Vliet, Roman Peter Jakob, Alberto Reinders, Alexander Klotz, Timm Maier, Urs Jenal

## Abstract

Cyclic diguanylate (c-di-GMP) is a ubiquitous second messenger that regulates a wide range of biological processes in bacteria, including motility, surface attachment, virulence and persistence. The regulatory networks controlling c-di-GMP are generally complex and understudied. This is largely due to a lack of appropriate tools to monitor dynamic changes of c-di-GMP concentrations *in vivo* in a non-invasive manner. Here, we develop a genetically-encoded ratiometric c-di-GMP biosensor, called cdGreen2, by applying a powerful directed evolution approach based on iterative fluorescence-activated cell sorting (FACS) under alternating c-di-GMP regimes. We demonstrate that cdGreen2 can robustly track c-di-GMP dynamics in live cells with ultra-high temporal resolution over multiple generations. To validate its exquisite diagnostic power, we utilize cdGreen2 to dissect the regulatory networks driving bimodal developmental programs in the environmental model organism *Caulobacter crescentus* and the human pathogen *Pseudomonas aeruginosa*. These studies disclose the molecular determinants governing cell cycle-dependent c-di-GMP oscillations in *C. crescentus* and surface-induced c-di-GMP asymmetry in *P. aeruginosa*. The sensitivity and versatility of cdGreen2 will help unveil c-di-GMP dynamics in a wide range of organisms with unprecedented temporal resolution. The simple, yet powerful design principles underlying cdGreen2 will serve as a blueprint for the development of similar, orthogonal biosensors for other signaling molecules, metabolites or antibiotics, paving the way to uncover the complex interplay of small molecule-based networks with unprecedented spatiotemporal resolution.

## INTRODUCTION

The ubiquitous second messenger c-di-GMP plays pivotal roles in many bacteria, regulating important behavioral and physiological processes like motility and virulence, or adherence to and growth on surfaces ^1–8^. Importantly, accurate control of c-di-GMP was shown to be critical for the establishment of infections and the development of resilience against host-mediated stress and antibiotic therapy in several human pathogens ^9^. In most bacteria, regulatory networks controlling c-di-GMP are complex with a multitude of enzymes being responsible for the controlled synthesis and degradation of the signaling compound ^10–12^. For example, the opportunistic human pathogen *Pseudomonas aeruginosa* encodes 38 enzymes involved in the «make and break» of c-di-GMP ^13^, and in some phyla enzymes regulating c-di-GMP constitute more than 1% of the organism’s total protein repertoire ^1^. This makes the genetic dissection of the respective regulatory pathways and mechanisms challenging. Much of the information on c-di-GMP control and its role in bacterial physiology and behavior stems from extrapolations of genetic data, generally complemented with invasive biochemical assays providing snapshots of c-di-GMP at steady-state levels in bulk populations. However, such measurements do not provide single cell information and thus fail to visualize signaling heterogeneity in bacterial populations, to track cell cycle-dependent fluctuations in asynchronous populations, or to distinguish distinct c-di-GMP-mediated cell fates in bacterial communities. Advancing our mechanistic understanding of c-di-GMP signaling in bacteria thus requires genetically encoded biosensors that monitor dynamic changes of c-di-GMP levels in real-time and in individual cells in a non-invasive manner.

The strong interest in the availability of robust biosensors for c-di-GMP or other small molecules has inspired repeated attempts to develop such tools. However, available biosensors generally suffer from major drawbacks. Transcription- or translation-based reporters, although readily available, are generally not suitable for real-time measurements due to considerable temporal delays caused by expression, folding, maturation or stability of the reporters ^14–19^. This limitation can be overcome by allosteric sensors operating on the posttranslational level. For instance, a recently described sensor based on bimolecular fluorescence complementation (BiFC) is suitable to measure steady-state levels of c-di-GMP ^20^. However, the irreversible nature of reconstituting a functional fluorescent protein by BiFC ^21^ precludes this tool from accurately measuring dynamic c-di-GMP fluctuations. More suitable biosensors that can potentially track changes in c-di-GMP levels in real time make use of Förster resonance energy transfer (FRET) between two fluorescent proteins sandwiching a c-di-GMP-binding protein ^22–24^, or of bioluminescence resonance energy transfer (BRET) where a ligand-binding protein is linked to a luciferase and a fluorescent protein ^25^. While FRET sensors can identify and monitor individual cells with different levels of c-di-GMP, they generally lack robustness and, so far, have not been shown to record dynamic changes of c-di-GMP over physiologically relevant time scales. In addition, FRET sensors generally suffer from limited dynamic ranges, making them highly susceptible to noise and imposing challenging downstream analysis procedures.

Given the lack of available tools to reliably report on c-di-GMP dynamics in real-time in individual cells, we set out to develop a novel biosensor that overcomes the above limitations and allows monitoring of c-di-GMP dynamics in a diverse range of bacteria by accessing several single-cell techniques, including live cell microscopy and flow cytometry. Starting from a scaffold comprising a circularly permuted enhanced green fluorescent protein (cpEGFP) sandwiched by c-di-GMP-binding domains, we applied a directed evolution-inspired approach ^26,27^, where c-di-GMP-responsive biosensors with increasing dynamic range and rapid binding kinetics were gradually selected by using iterative fluorescence-activated cell sorting (FACS) under alternating c-di-GMP regimes. The resulting biosensor, termed cdGreen2, was benchmarked by dissecting the c-di-GMP regulatory networks of two model organisms, the environmental bacterium *Caulobacter crescentus* and the human pathogen *Pseudomonas aeruginosa*. The bimodal developmental program of *C. crescentus* generates a motile and planktonic swarmer (SW) cell and a sessile, surface-attached stalked (ST) cell. Cell differentiation was proposed to depend on precise and genetically hardwired oscillations of c-di-GMP that coordinate cell cycle progression with morphogenesis and behavior ^3,19^. Similarly, *P. aeruginosa* undergoes a surface-induced asymmetric program generating motile and sessile offspring to maximize surface colonization ^4,28^. Like in *C. crescentus*, this program was postulated to be orchestrated by the asymmetric distribution of c-di-GMP during the initial cell divisions of surface adherent *P. aeruginosa* cells. Using cdGreen2 to visualize c-di-GMP in individual cells, we largely corroborate these models and define the molecular mechanisms underlying the bimodal phenotypic specialization. We anticipate that cdGreen2 will become a standard molecular tool for the scientific community to dissect the function and mechanisms of c-di-GMP in a large variety of bacteria.

## RESULTS AND DISCUSSION

### Design and directed evolution of a biosensor for c-di-GMP

To develop a biosensor for c-di-GMP, we took advantage of the observation that the optical properties of the chromophore in GFP-derived fluorescent proteins are highly sensitive to the immediate microenvironment ^29–33^. We selected a circularly permutated EGFP (cpEGFP) where the N- and C-termini were engineered to be located close to the chromophore ^34^ and fused the C-terminal c-di-GMP binding domain (CTD) of the transcription factor BldD (BldD^CTD^) ^35^ to both termini (Fig. 1a). BldD is monomeric in the absence of c-di-GMP, but homo-dimerizes in the presence of the ligand with two intercalated dimers of c-di-GMP bridging the BldD momomers (Fig. 1b,c). We rationalized that such fusion proteins could change fluorescence intensity upon c-di-GMP binding due to local perturbations around the chromophore. Because we presumed that maximal biosensor functionality would require optimizing the length and flexibility of the linkers connecting cpEGFP with the BldD protomers, we constructed a library of the cpEGFP-[BldD^CTD^]_2_ fusion protein with randomized linker length and amino acid composition. This library was then transferred to *Escherichia coli* “cdG tuner” strain AKS494 (Fig. 1d), in which the concentration of c-di-GMP could be adjusted from <50 nM to >5 µM ^36^. Iterative fluorescence-activated cell sorting (FACS) under alternating c-di-GMP high and low regimes was then used to enrich for functional biosensors displaying maximal differences in fluorescence intensity between the two c-di-GMP states (Fig. 1d). This led to the isolation of a first-generation biosensor with an approximately three-fold change in fluorescent intensity when expressed in *E. coli* cells with low and high c-di-GMP concentrations, respectively (Fig. 1e). Sequential cycles of linker optimization and selection (see Materials and Methods) led to gradually improved versions of the biosensor, generating a final variant termed cdGreen with a more than ten-fold change in fluorescent intensity between high and low c-di-GMP states (Fig. 1e).

**Figure 1.**
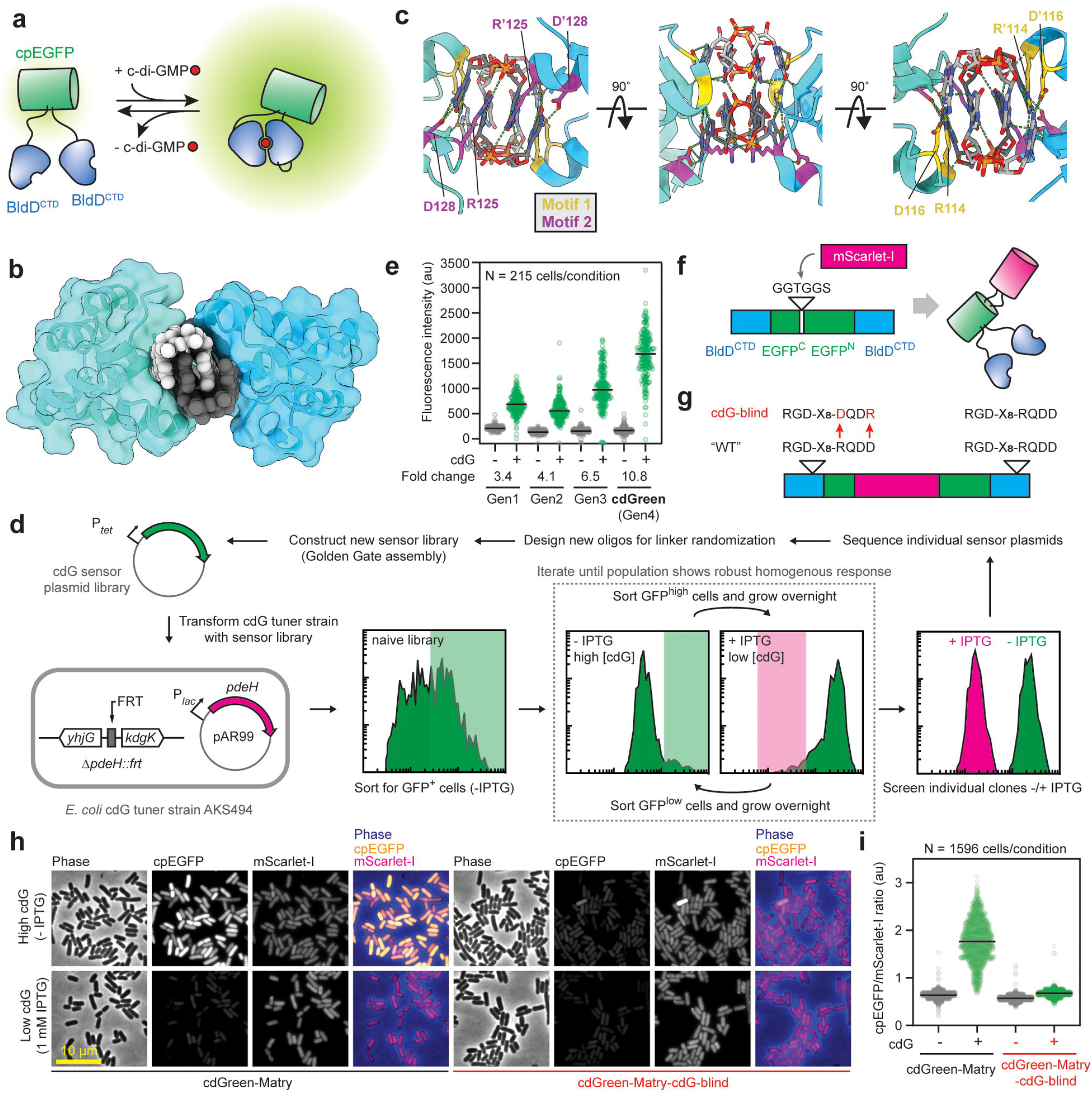
Design and isolation of c-di-GMP biosensor cdGreen. a. Schematic of cdGreen biosensor design and function. Upon c-di-GMP-mediated dimerization of the two BldD^CTD^ protomers, the cpEGFP moiety undergoes a conformational change resulting in increased fluorescent intensity. b. Cartoon representation of the dimeric BldD^CTD^ in complex with tetrameric c-di-GMP (PDB: 5KHD). The intercalated c-di-GMP dimer binding to motif 1 is shown with individual c-di-GMP molecules in light and dark grey, respectively. Note that dimerization is solely mediated by c-di-GMP without any contribution from protein-protein interactions. c. Close-up of the BldD^CTD^-c-di-GMP interface with key residues (underlined) in motif 1 (<u>R</u>G<u>D</u>) and motif 2 (<u>R</u>QD<u>D</u>) highlighted as yellow and purple sticks, respectively. Based on PDB: 5KHD. d. Schematic outline of the iterative FACS approach used to isolate c-di-GMP biosensors, including the 4^th^ generation biosensor cdGreen (see Material and Methods for details). e. Performance of different c-di-GMP biosensors in c-di-GMP high and low conditions assayed in strain AKS494 by microscopy. For “c-di-GMP low” conditions, cells were grown with 1 mM IPTG. 200 nM aTc was included in all conditions for expression of the biosensors. For image analysis, images were background-corrected. f. Schematic of the cdGreen-Matry construct. g. Schematic of the cdG-blind cdGreen-Matry construct. h. Microscopy snapshots of AKS494 harboring plasmids p2H12-Matry or p2H12-Matry-cdG-blind grown with (1 mM) or without IPTG. 200 nM aTc was included in all conditions for expression of the biosensors. i. Quantification of c-di-GMP levels (cpEGFP/ mScarlet-I ratio) of strains shown in panel f.

Because single fluorescent protein biosensors (SFPBs) are intensiometric by design and their readout prone to naturally occurring variations of protein concentration *in vivo*, we sought to normalize c-di-GMP-mediated changes in fluorescence intensity by fusing the sensor to a reference fluorescent protein (FP). To avoid interference of the reference FP with biosensor functionality, we followed the recently developed “Matryoshka” approach ^37^ and inserted the mScarlet-I ^38^ reference FP in the loop connecting the original EGFP N-and C-termini (Fig. 1f,g). The resulting construct, termed cdGreen-Matry, retained its c-di-GMP sensitivity *in vivo* (Fig. 1h) and made it possible to normalize the c-di-GMP (cpEGFP) signal with the reference FP (Fig. 1i), thus effectively converting the initial intensiometric into a ratiometric biosensor.

### cdGreen is a ratiometric c-di-GMP biosensor with high ligand specificity and dynamic range

To demonstrate that the biosensor specifically responds to c-di-GMP, we interchanged the Arg and Asp residues of ligand-binding motif 2 in one of the BldD^CTD^ protomers (Fig. 1c,g). These substitutions were previously shown to fully abrogate c-di-GMP binding in the context of full-length BldD ^35,39^. Accordingly, the mutant sensor failed to respond to c-di-GMP and displayed signals in agreement with the c-di-GMP state (Fig. 1h,i). To characterize our biosensors in more detail, we expressed and purified hexa-His tagged cdGreen and cdGreen-Matry to homogeneity (Supplementary Fig. 1). Purified cdGreen showed two major excitation peaks at 497 nm and 405 nm with corresponding emission maxima at 513 and 518 nm, respectively (Fig. 2a). Importantly, the addition of c-di-GMP strongly increased the excitation peak at 497 nm, but reduced the peak at 405 nm in a dose-dependent manner with concomitant changes of the corresponding emission peaks (Fig. 2a,b). This inverse response of two major excitation peaks to c-di-GMP provides a ratiometric readout, a feature that is critical for *in vivo* applications as it provides an elegant means to internally normalize stochastic cell-to-cell variations of biosensor concentrations. cdGreen-Matry showed a similar c-di-GMP-dependent behavior, but a reduced response amplitude due to Förster resonance energy transfer (FRET) between its cpEGFP and mScarlet-I moieties (Fig. 2a). The dose-response curves recorded for cdGreen upon excitation at 497 nm and 405 nm and emission at 530 nm revealed sigmoidal monotonic behavior for both wavelengths with maximal fold changes of 12.0 and 5.0 and fitted K_d_s of 214 nM (95% CI: 210-217 nM; Hill slope = 2.30) and 225 nM (95% CI: 217-228 nM; Hill slope = -2.43), respectively (Fig. 2c). The combined ratiometric readout also displayed a sigmoidal and monotonic behavior with a dynamic range of almost 70 and a fitted K_d_ of 386 nM (95% CI: 381-391 nM; Hill slope: 3.31) (Fig. 2c). The response of purified cdGreen was specific to c-di-GMP, as none of the related nucleotides tested induced a response or caused interference with the ability of cdGreen to respond to c-di-GMP (Supplementary Fig. 2a). Stoichiometric titration experiments revealed a plateau at a ratio of 2 c-di-GMP molecules per 1 molecule of protein (Supplementary Fig. 2b). This is in line with the observation that BldD, although binding two intercalated c-di-GMP dimers in the fully functional state (Fig. 1b,c), is able to dimerize with only one c-di-GMP dimer bound ^39^. Isothermal titration calorimetry (ITC) experiments showed two consecutive binding events of c-di-GMP to cdGreen, likely corresponding to dimeric and tetrameric c-di-GMP (Supplementary Fig. 2c). These results demonstrated that cdGreen binds tetrameric c-di-GMP, but that one intercalated c-di-GMP dimer is sufficient to provoke the maximal biosensor response.

**Figure 2.**
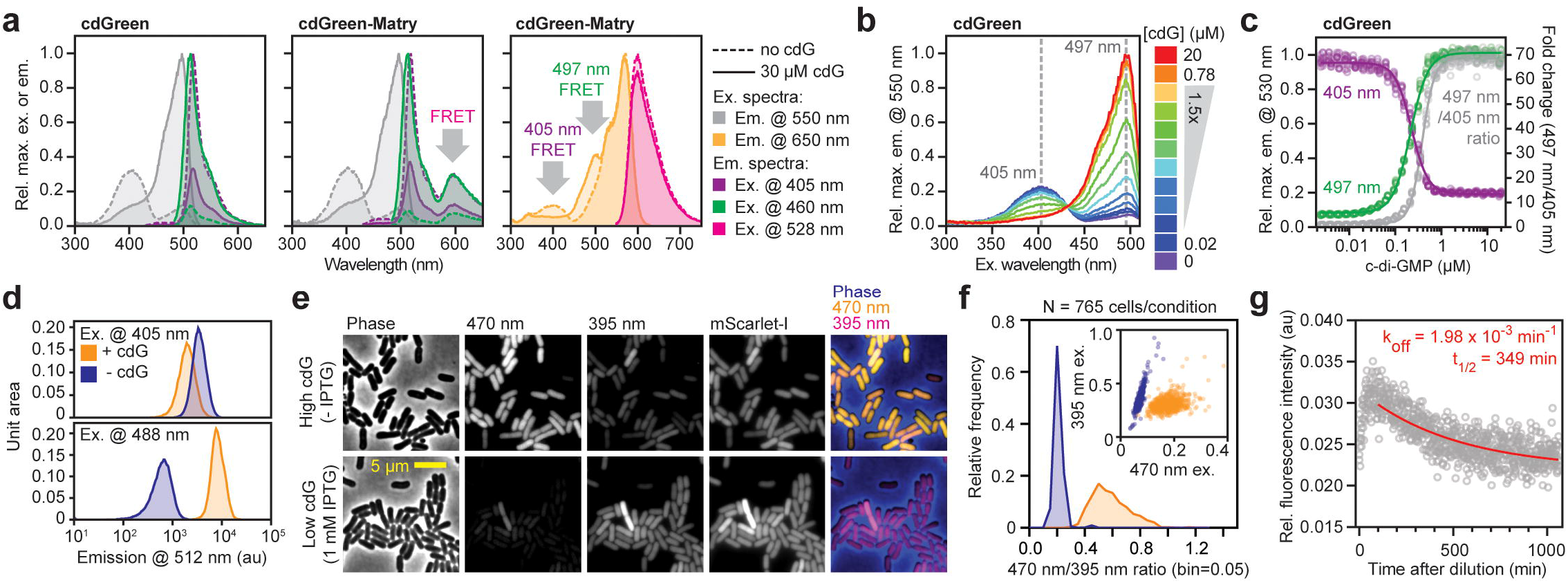
*In vitro* characterization of c-di-GMP biosensors cdGreen and cdGreen-Matry. a. Spectral excitation and emission scans of cdGreen and cdGreen-Matry. Peaks resulting from FRET between cpEGFP and mScarlet-I in cdGreen-Matry are indicated. b. Excitation spectra of cdGreen with different c-di-GMP concentrations. c. C-di-GMP dose-response curve of cdGreen emission at 530 nm with excitation at 405 nm and 497 nm (left y-axis) and the combined ratiometric readout (ratio of 550 nm emission signal with 497 nm and 405 nm; right y-axis). d. Flow cytometry of strain AKS494 harboring plasmid p2H12 grown with (1 mM) or without IPTG. 200 nM aTc was included in all conditions for expression of the biosensors. e. Microscopy images of strain AKS494 carrying plasmid pConRef-2H12 grown with (1 mM) or without IPTG. f. Quantification of c-di-GMP levels (470 nm/ 395 nm ratio) of strains shown in panel e. g. Dissociation kinetics of cdGreen as determined by the “dissociation by dilution” method (see Material and Methods for details).

To test whether the ratiometric biosensor readout of c-di-GMP could also be used *in vivo*, we next expressed cdGreen in the *E. coli* “cdG tuner” strain AKS494 (Fig. 1d) in c-di-GMP high and low conditions. Using flow cytometry (Fig. 2d) or live-cell imaging (Fig. 2e,f), signals were readily detected in the GFP/FITC emission channel upon excitation at 395-405 nm, with signal strengths anti-correlating with the GFP/FITC emission channel signals upon excitation with the standard GFP/FITC excitation wavelengths (470-488 nm). Thus, the ratio of emissions upon excitation with the two major excitation wavelengths can be used for sensor normalization *in vivo*. Excitation around 400 nm is routinely used for DAPI excitation and thus is a standard setting in most modern fluorescence microscopes and flow cytometry instruments, thus allowing the direct adoption of our novel c-di-GMP biosensor in many standard laboratory settings without the need of specialized equipment.

### Isolation of cdGreen2, a dynamic and robust ratiometric c-di-GMP biosensor with rapid off kinetics

To evaluate the potential of cdGreen to read out dynamic changes of c-di-GMP *in vivo*, we next determined the dissociation rate constant (k_off_) using the “dissociation by dilution” method ^40^ (see Materials and Methods for details). This revealed a k_off_ of 1.98 × 10-3 min-1 (95% CI: 1.62-2.36 × 10^−3^ min^-1^) corresponding to a half-life of 349 min (95% CI: 294-428 min) (Fig. 2g). Based on the experimentally determined k_off_ and K_d_ values, we calculated an association rate constant (k_on_ = k_off_ K_d_ ^-1^) of around 9000 M^-1^ min^-1^. Thus, although the dynamic range of cdGreen is exceptionally high, slow dissociation of c-di-GMP likely precludes dynamic measurements *in vivo*. To test this notion, we expressed cdGreen in *Caulobacter crescentus*, an aquatic bacterium with a characteristic asymmetric division cycle that generates motile and sessile offspring. While levels of c-di-GMP are high in sessile stalked cells (ST), newborn swarmer cells (SW) experience a short trough of c-di-GMP during their motile phase, before c-di-GMP levels sharply rise again to drive morphogenesis back into sessile replication-competent ST cells (Fig. 3a) ^41,42^. In line with the k_off_ determined *in vitro*, cdGreen expressed in *C. crescentus* reported constant high levels of c-di-GMP and failed to reproduce its expected fluctuation during the cell cycle (Supplementary Fig. 3a). A signal corresponding to low c-di-GMP levels was observed when cdGreen was expressed in a *C. crescentus* mutant unable to produce this second messenger (NA1000 rcdG^0^) ^41^ (Supplementary Fig. 3b). This suggested that, while cdGreen is fully functional in *C. crescentus*, it fails to report on the expected dynamic fluctuations of the second messenger, likely due to slow dissociation kinetics.

**Figure 3.**
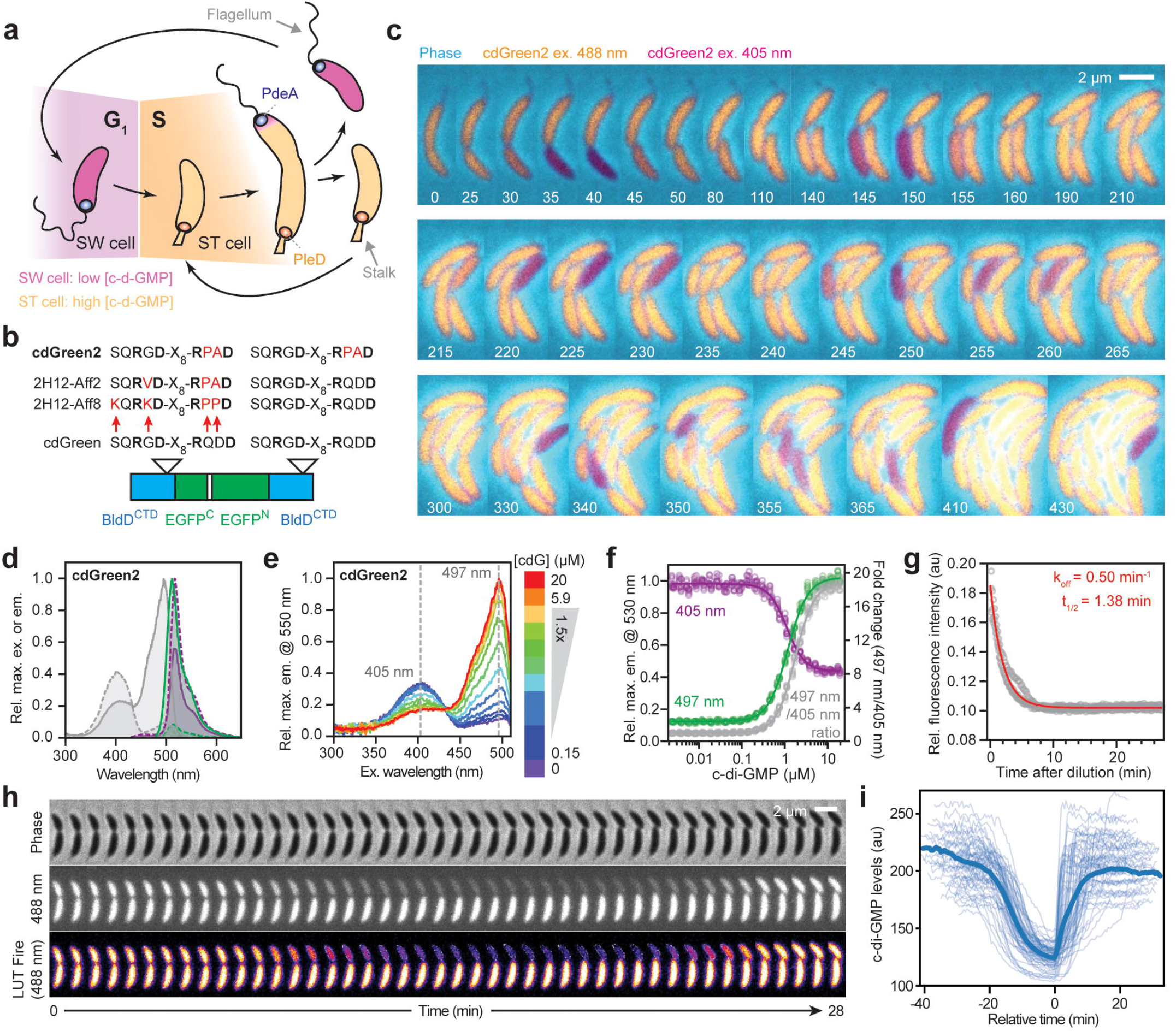
Isolation and *in vitro* characterization of c-di-GMP biosensor cdGreen2. a. Schematic of c-di-GMP oscillations during the *C. crescentus* cell cycle. Stalk and flagellum are indicated, as well as the opposite polar localization of the major diguanylate cyclase PleD at the stalked pole and the major phosphodiesterase PdeA at the incipient flagellated pole. b. Schematic of amino acid changes in cdGreen derivatives 2H12-Aff2, 2H12-Aff8 and cdGreen2. c. Time-lapse series of strain NA1000 carrying plasmid pQFmcs-2H12.D11. Time stamps indicate minutes. Images are overlays of the cdGreen2 FITC/FITC signal pseudo-colored in yellow, the cdGreen2 DAPI/FITC signal pseudo-colored in magenta and the phase contrast image pseudo-colored in cyan. Also see related Supplementary Fig. 4d. d. Spectral excitation and emission scans of cdGreen2. Emission and excitation wavelengths are color-coded as in Fig. 2a. e. Excitation spectra of cdGreen2 with different c-di-GMP concentrations. f. C-di-GMP dose-response curve of cdGreen2 emission at 530 nm with excitation at 405 nm and 497 nm (left y-axis) and the combined ratiometric readout (ratio of 550 nm emission signal with 497 nm and 405 nm; right y-axis). g. Dissociation kinetics of cdGreen2 as determined by the “dissociation by dilution” method (for details see Material and Methods). h. High-temporal-resolution imaging of c-di-GMP levels (20-sec intervals) during *C. crescentus* division and G1-S phase transition. Note that only every second frame is shown. i. Quantification of single-cell c-di-GMP dynamics in multiple cells. Individual cell tracks were overlaid setting the time point at which a cell showed minimum c-di-GMP levels as a reference point (t=0).

Because the affinity equilibrium dissociation constant K_d_ directly relates to the dissociation and association rate constants k_off_ and k_on_ (via K_d_ = k_off_ k_on_ ^-1^), we reasoned that variants with lower binding affinities may show more rapid off kinetics. We thus generated mutant libraries with randomized amino acids at non-conserved positions in the immediate vicinity of residues involved in c-di-GMP binding (Fig. 1c,3b) and subjected them to iterative FACS-based enrichment for variants that shift to the off state at higher c-di-GMP concentrations as compared to cdGreen (see Materials and Methods). This gave rise to two variants, 2H12-Aff2 and 2H12-Aff8, both of which had Pro residues in position 2 of the RXXD ligand binding motif (Fig. 1c,g; Fig. 3b). We speculate that the Gln126Pro substitution repositions the key ligand-binding residues R125 and D128 in a way that c-di-GMP is bound less tightly compared to parental cdGreen. Because 2H12-Aff2 and 2H12-Aff8 failed to recapitulate expected c-di-GMP oscillations in *C. crescentus* (Fig. 3a; Supplementary Fig. 4a), we constructed additional cdGreen-based derivatives by combining substitutions isolated in 2H12-Aff2 and 2H12-Aff8 variants in one or both BldD^CTD^ protomers. Variants were scored for their ability to visualize the anticipated differences in c-di-GMP concentration in *C. crescentus* ST and SW cells (Fig. 3a). This approach revealed a single sensor variant – termed cdGreen2 – harboring a RPAD motif 2 variant in both BldD^CTD^ protomers that recapitulated the expected c-di-GMP oscillations over the *Caulobacter* cell cycle (Fig. 3c; Supplementary Fig. 4a,b). Cells expressing cdGreen2 displayed constantly high levels of c-di-GMP in their ST phase, while levels dropped in newborn SW cells for a period of 20-30 minutes, before reaching their original levels upon SW-to-ST cell differentiation (Fig. 3c). Co-expression of cdGreen2 with a ST cell-specific marker (SpmX-mCherry) ^19,43^ confirmed that the drop in c-di-GMP coincided with the motile SW stage of the *Caulobacter* cell cycle (Supplementary Fig. 4c). Importantly, cdGreen2 allowed robust imaging of the expected cell cycle-dependent c-di-GMP fluctuations over several hours and generations without an apparent decrease in sensor performance (Fig. 3c; Supplementary Fig. 4d).

**Figure 4.**
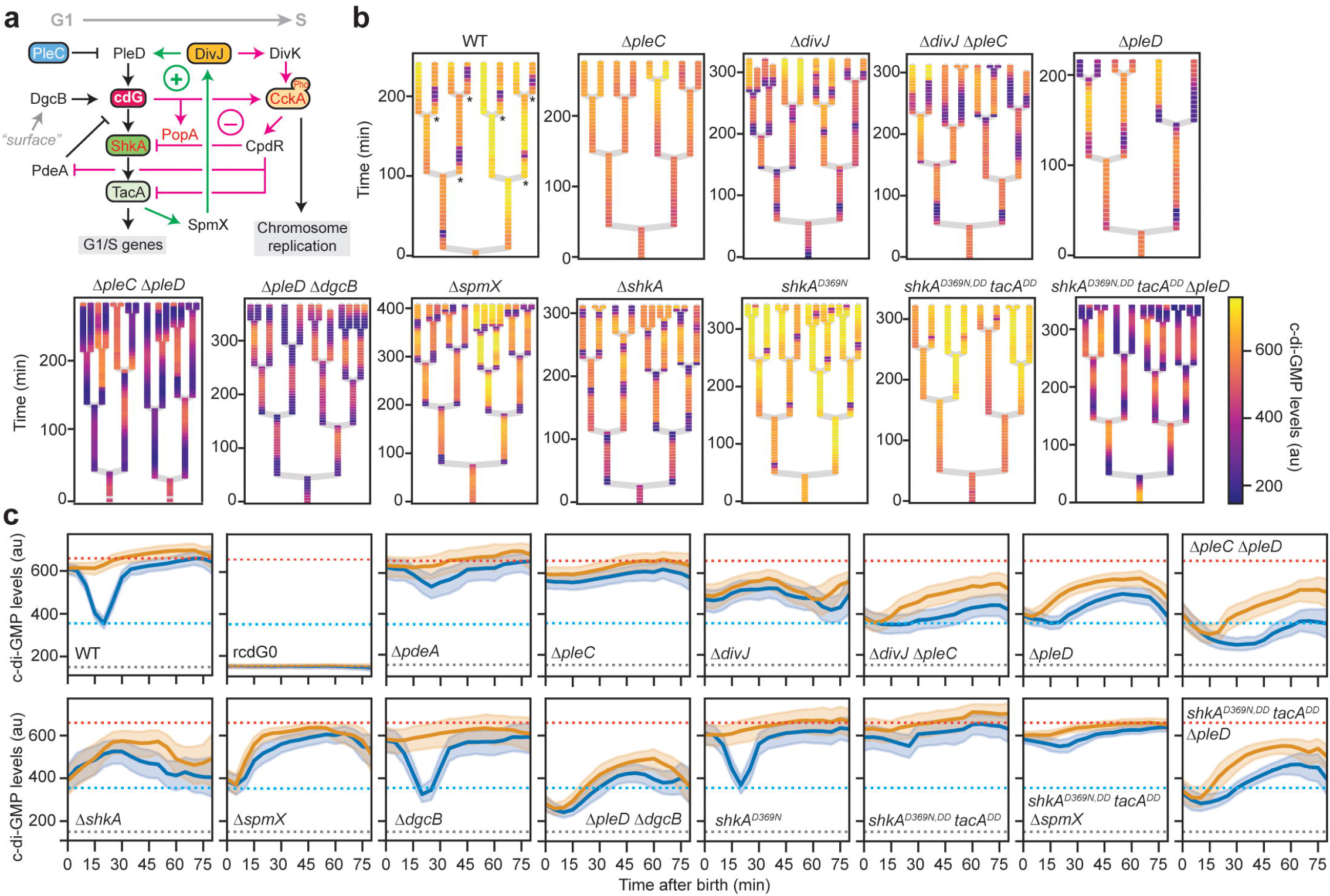
Dissecting the G1-S-specific c-di-GMP network of *C. crescentus*. a. Schematic of the regulatory network driving G1-S transition in *C. crescentus*. C-di-GMP effector proteins, i.e., proteins whose functions are directly controlled by c-di-GMP binding, are highlighted in red. CpdR and PopA are ClpX-specific adapter proteins required for ShkA, TacA and PdeA proteolysis; PopA activity requires c-di-GMP binding. The positive and negative feedback loops (see main text for details) during G1-S phase transition are highlighted by green and magenta arrows, respectively. b. Lineage trees of indicated strains carrying plasmid pQFmcs-2H12.D11-scarREF. Asterisks in the “wild type” panel indicate SW cells; note that ST cells show shorter interdivision times than SW cells. c. Tracking of c-di-GMP levels in pairs of sister cells after cell division. For details, see Material and Methods. The orange dotted line indicates average c-di-GMP levels in wild-type ST cells, the blue dotted line indicates average minimal c-di-GMP levels in wild-type SW cells; the grey dotted line indicates average c-di-GMP levels in rcdG^0^ cells and thus reflects the background signal in absence of c-di-GMP.

Purified cdGreen2 showed spectral properties similar to parental cdGreen with two major excitation peaks at 497 nm and 405 nm with corresponding emission maxima at 513 and 518 nm, respectively (Fig. 3d). Addition of c-di-GMP increased the emission at 550 nm upon 497 nm excitation and concomitantly decreased the emission at 550 nm upon 405 nm excitation in a dose-dependent manner (Fig. 3d,e). Dose-response curves at 497 nm and 405 nm excitation with emission at 530 nm showed sigmoidal monotonic behavior with maximal fold changes of 8.2 and 2.2 and with fitted K_d_s of 1.24 µM (95% CI: 1.22-1.26 µM; Hill slope = 1.86) and 1.17 µM (95% CI: 1.12-1.21 µM; Hill slope = -2.07), respectively. The combined ratiometric readout displayed a sigmoidal and monotonic behavior with a maximal fold change of almost 20 and a fitted K_d_ of 1.83 µM (95% CI: 1.81-1.85 µM; Hill slope: 1.98) (Fig. 3f). Stoichiometric titration experiments revealed a 2:1 ligand-to-protein ratio, similar to parental cdGreen (Supplementary Fig. 2d). Finally, “dissociation by dilution” experiments ^40^ (see Materials and Methods) with purified cdGreen2 determined a dissociation rate constant (k_off_) of 0.50 min^-1^ (95% CI: 0.49-0.52 min^-1^) corresponding to a half-life of 1.38 min (95% CI: 1.33-1.43 min) (Fig. 3g). The calculated association rate constant (k_on_ = k_off_ K_d_ ^-1^) is around 400’000 M^-1^ min^-1^. These kinetic values were largely confirmed by independent experiments globally fitting parameters based on association kinetics in the pseudo-first-order regime (see Materials and Methods) (Supplementary Fig. 2e,f).

In line with the above, time-lapse microscopy of growing *C. crescentus* cells expressing cdGreen2 at high temporal resolution (20-sec intervals) exposed a robust c-di-GMP trough period of about 20-30 minutes in differentiating SW cells (Fig. 3h,i). Importantly, no photo-toxicity or bleaching effects were observed when cells were imaged with high temporal resolution for up to 24 hours (Supplementary Movie 1). This exceptional robustness likely relates to the overall signal strength of cdGreen2, allowing the use of ultra-short exposure in the fluorescent channels and reduced laser power during time lapse experiments. In sum, these results demonstrate that cdGreen2 affords robust, quantitative and dynamic monitoring of c-di-GMP at the single cell level with ultra-high temporal resolution and without phototoxic effects or photo-bleaching over multiple generations.

### cdGreen2 unveils the regulatory network driving *Caulobacter* morphogenesis

Accurate temporal and spatial control of c-di-GMP was proposed to be a central element of *C. crescentus* morphogenesis and cell cycle progression. Synthesis and degradation of c-di-GMP are controlled by two key enzymes, the diguanylate cyclase PleD and the phosphodiesterase PdeA, which position to opposite poles of dividing cells and differentially partition into ST and SW progeny during division ^3^ (Fig. 3a, 4a). We used cdGreen2 to investigate the spatial and temporal control of these enzymes in detail. C-di-GMP asymmetry and oscillation were visualized over multiple generations in lineage trees originating from single dividing cells expressing cdGreen2 (Fig. 4b). Analyzing multiple sister pairs after division in the wild type (n=108) emphasized the highly deterministic nature of c-di-GMP asymmetry during the bimodal *Caulobacter* cell cycle (Fig. 4c). Establishing c-di-GMP asymmetry requires the complete separation of the cellular compartments during cytokinesis. Cells depleted for the essential division protein FtsZ by means of CRISPRi-mediated FtsZ depletion failed to constrict, resulting in filamentous cells that maintained high levels of c-di-GMP throughout the course of the experiment (Supplementary Fig. 5a). A few rare cells that managed to escape FtsZ depletion, initiated cytokinesis and rapidly established c-di-GMP asymmetry (Supplementary Fig. 5b). Thus, the diguanylate cyclase PleD is dominant over PdeA when positioned in the same cellular compartment, with different cell fates ultimately being enforced by a checkpoint during cytokinesis.

**Figure 5.**
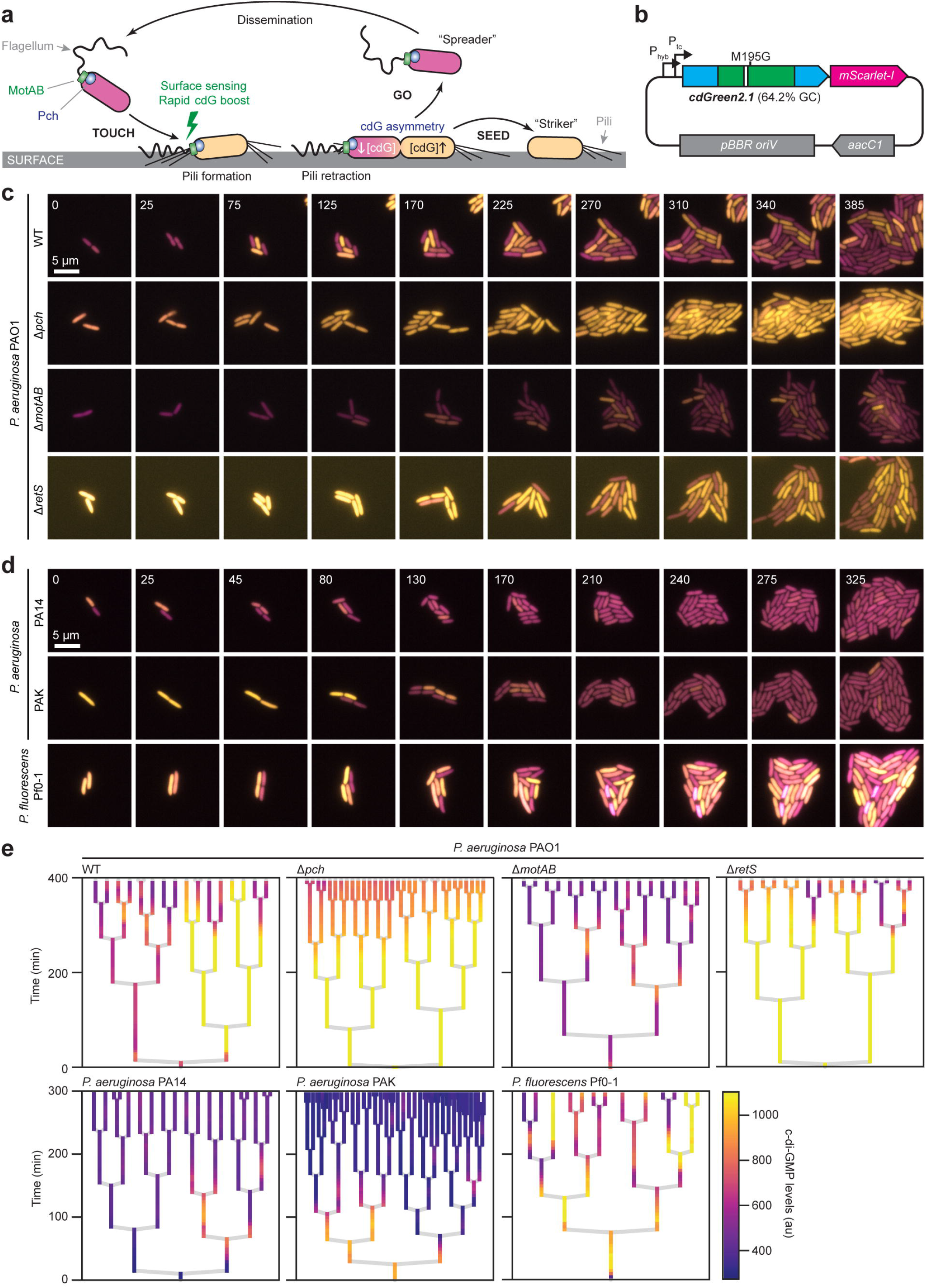
C-di-GMP dynamics in surface-associated *Pseudomonas*. a. Schematic of the “touch-seed-and-go” model in *P. aeruginosa* PAO1. b. Schematic of the plasmid used in *Pseudomonas* expressing cdGreen2.1 and the reference FP mScarlet-I. Note that cdGreen2.1, besides being high-GC-content optimized, also contains a single codon change (Met195Gly) that removes a potential internal start codon. P_hyb_ and P_tc_ represent two custom-designed nested, constitutive promoters. c. Tracking of c-di-GMP levels in indicated *P. aeruginosa* PAO1 strains after spotting on agarose pads. Images are overlays of the cdGreen2.1 FITC/FITC signal pseudo-colored in yellow and the mScarlet-I signal pseudo-colored in magenta. Time stamps indicate minutes. Contrast and brightness settings are the same for all images. d. Tracking of c-di-GMP levels in indicated *P. aeruginosa* strains after spotting on agarose pads. Images are overlays of the cdGreen2.1 FITC/FITC signal pseudo-colored in yellow and the mScarlet-I signal pseudo-colored in magenta. Time stamps indicate minutes. Contrast and brightness settings are the same for all images. e. Lineage trees of indicated strains.

Genetic evidence had suggested that levels of c-di-GMP are kept low in SW cells by PdeA ^44^ and by the phosphatase PleC, which retains PleD in its inactive, unphosphorylated state ^45–48^ (Fig. 4a). In line with this, c-di-GMP failed to drop in newborn SW cells of mutants lacking PdeA or PleC (Fig. 4b,c; Supplementary Fig. 5c,d). Deleting *pleD* in the Δ*pleC* mutant lowered levels of c-di-GMP and essentially phenocopied a Δ*pleD* single mutant (see below), demonstrating that PleD is indeed over-activated in the absence of the SW cell-specific phosphatase PleC (Fig. 4b,c). Levels of c-di-GMP were significantly reduced in a Δ*pleD* mutant, and were even lower in a strain that also lacked DgcB, a second important diguanylate cyclase, which was recently shown to promote rapid surface attachment of SW cells in response to mechanical cues ^49^. Although a Δ*pleD* Δ*dgcB* double mutant showed strongly reduced levels of c-di-GMP levels, the concentration of c-di-GMP still changed over the cell cycle, indicating that additional, minor diguanylate cyclases are involved in this process ^41^. However, the observation that c-di-GMP levels in SW and ST cell progeny were indistinguishable in the double mutant (Fig. 4b) indicated that PleD and DgcB are the principal drivers of *Caulobacter* asymmetry.

As *Caulobacter* cells transit from G1 into S phase, c-di-GMP levels gradually increase ^41,42^ after PleD is activated by phosphorylation ^48^ and PdeA is proteolytically removed ^44^. This leads to a series of accurately timed events prompting exit from G1, SW-to-ST cell morphogenesis and entry into S phase. First, c-di-GMP-dependent activation of the ShkA-TacA pathway stimulates the expression of hundreds of genes that orchestrate the morphological restructuring of the motile SW cells into sessile ST cell ^19,50–52^. This sets in motion a positive feedback loop that includes the kinase DivJ and its activator SpmX, leading to the production of more c-di-GMP via the reinforced activation of PleD (Fig. 4a). The resulting buildup of c-di-GMP was proposed to trigger a second, negative feedback loop via the cell cycle kinase CckA that terminates ShkA-mediated gene expression and initiates chromosome replication upon entry into S-phase ^19,50,53–56^. While this complex regulatory network was proposed to accurately time *Caulobacter* development (Fig. 4a), direct visualization of c-di-GMP in support of the model is missing. We now show that mutants lacking components of the proposed positive feedback loop (ShkA, SpmX, DivJ) all showed reduced c-di-GMP levels. For example, a Δ*divJ* mutant failed to increase the concentration of c-di-GMP during G1-S transition and, similar to a Δ*pleD* mutant, showed strongly compromised c-di-GMP asymmetry during cell division (Fig. 4b,c). A Δ*divJ* Δ*pleC* double mutant largely phenocopied a Δ*divJ* mutant with respect to c-di-GMP dynamics, suggesting that PleD remains mostly unphosphorylated and inactive in this background. Mutants lacking ShkA or SpmX showed similar phenotypes, supporting their proposed role in robustly boosting c-di-GMP levels during the G1-S transition (Fig. 4b,c). Interestingly, Δ*pleD*, Δ*divJ*, Δ*spmX* and Δ*shkA* single mutants, as well as the Δ*divJ* Δ*pleC* double mutant, showed a premature trough of c-di-GMP in late predivisional cells that then extended into both daughter cells after division. This observation indicated that (i) PdeA and possibly other phosphodiesterases are active already before cell constriction, and that (ii) robust activation of PleD is important to overrule the phosphodiesterase(s) and to maintain the identity of ST cells.

Interference with components of the negative feedback loop had the opposite effect. Introducing amino acid substitutions in ShkA and TacA that protected them from being degraded (*shkA*^*DD*^ *tacA*^*DD*^) ^19,57^ in combination with a mutation that rendered ShkA constitutively active (*shkA*^*D369N*^) ^19^ led to constitutively high levels of c-di-GMP, bypassing the characteristic drop of c-di-GMP in SW cells (Fig. 4b,c). Deletion of *pleD* in the *shkA*^*D369N,DD*^ *tacA*^*DD*^ background essentially phenocopied a Δ*pleD* mutant, confirming that the negative feedback loop mediating ShkA and TacA proteolysis limits PleD-mediated activation of the ShkA-TacA pathway to a short time window, thereby providing accurate gene expression control during the G1-S transition. Deletion of *spmX* in the *shkA*^*D369N,DD*^ *tacA*^*DD*^ background had no such effect, suggesting that other direct or indirect ShkA/TacA-dependent factors stimulate PleD activity. Together, these results provide strong support for a developmental model in which the gradual upshift of c-di-GMP and the precise timing of gene expression during the *C. crescentus* cell cycle is executed by consecutive positive and negative feedback loops. Our findings also demonstrate that accurate control of the diguanylate cyclase PleD is of key importance to establish c-di-GMP asymmetry in *C. crescentus* ^41,55^. Finally, these experiments provide a convincing example for the potential of the cdGreen2 biosensor as a powerful cellular tool to directly correlate cellular dynamics and specific gene functions with changes of a small signaling molecule in individual cells.

### Recording c-di-GMP during *Pseudomonas aeruginosa* surface colonization

To expand the use of the cdGreen biosensor to pathogenic bacteria, we selected *Pseudomonas aeruginosa*, an opportunistic human pathogen that is notorious for its capacity to colonize and breach mucosal linings and cause acute and chronic infections. We have recently uncovered a sophisticated asymmetric program termed “touch-seed-and-go” that allows *P. aeruginosa* to effectively colonize surfaces ^28^. We had proposed that upon surface contact *P. aeruginosa* senses mechanical cues with its polar rotary flagellar motor and, in response, triggers a rapid upshift of c-di-GMP to stimulate surface attachment via the production of adhesive Type IV pili ^58^. Surface attached cells then undergo an asymmetric division, generating an offspring that retains high levels of c-di-GMP and remains surface-associated, and an offspring that reduces its internal c-di-GMP to regain motility and to explore more distant sites ^28^ (Fig. 5a). The reduction of c-di-GMP in one offspring relies on the activity of the phosphodiesterase Pch, which associates with the chemotaxis machinery at the flagellated pole ^59^. Using a codon-adapted version of cdGreen2, called cdGreen2.1, we scrutinized the proposed *P. aeruginosa* surface colonization model. Because autofluorescence in the violet excitation range under certain growth conditions of *P. aeruginosa* complicates internal normalization using the 405 nm excitation peak, we additionally included a transcriptionally coupled reference FP (mScarlet-I) in the construct expressing cdGreen2.1 for normalization (Fig. 5b).

Using cdGreen2.1, c-di-GMP was readily detected in *P. aeruginosa* strain PAO1 grown on LB agarose pads. As predicted by the “touch-seed-and-go” model, the first few division events on surface generated offspring with asymmetric c-di-GMP distributions (Fig. 5c). This was followed by a gradual increase of c-di-GMP heterogeneity with an increasing fraction of newborn cells adopting a low c-di-GMP state, arguing that genetic and environmental factors gradually influence the c-di-GMP program upon surface colonization (Fig. 5b). A strain lacking the phosphodiesterase Pch had lost its asymmetry, displaying uniformly high concentrations of c-di-GMP for an extended period (Fig. 5b,d). In line with the proposed role of the polar flagellum in sensing mechanical cues upon surface contact ^49,28^, a strain lacking the motor components MotA and MotB failed to increase c-di-GMP levels throughout the first few divisions on surface. Cell-to-cell variability of c-di-GMP was still observed in this strain, although at strongly reduced levels, indicating that pathways independent of flagellar-based mechanosensation contribute to an upshift of c-di-GMP under these conditions (Fig. 5a,c). Finally, we tested the role of RetS in c-di-GMP control, an atypical histidine kinase in the Gac/Rsm pathway ^60–62^ that was reported to negatively influence c-di-GMP levels in the *P. aeruginosa* strain PAK ^7^. Indeed, a Δ*retS* mutant of strain PAO1 showed strongly increased levels of c-di-GMP even at very early time points of surface exposure. This and the observation that the characteristic asymmetric distribution of c-di-GMP was strongly delayed in this mutant (Fig. 5b,d), argued that the role of the Gac/Rsm pathway is to limit c-di-GMP in planktonic cells of *P. aeruginosa* to provide them with the necessary sensitivity to respond to surface encounters by rapidly boosting c-di-GMP levels leading to effective surface adherence via c-di-GMP-mediated Type IV pili assembly ^28^. While the surface-mediated boost and subsequent asymmetry of c-di-GMP was highly pronounced in *P. fluorescens* Pf0-1, an environmental *Pseudomonas* isolate, *P. aeruginosa* strains PA14 and PAK showed only weak responses to surface exposure (Fig. 5d,e), indicating that they adapt to surfaces differently than strain PAO1. Given the observation that c-di-GMP often counters virulence of *P. aeruginosa* and other bacterial pathogens ^1^, it is possible that differential c-di-GMP control explains the hyper-virulent phenotype of strain PA14 ^63,64^. In this context, it is worth noting that the hyper-virulence of strain PA14 has previously been linked to a mutation in *ladS*, affecting Gac/Rsm pathway activity ^65^.

Altogether, these examples demonstrate the versatility and adaptability of the cdGreen2 biosensor for the detailed analysis of a diverse range of bacteria. Its ability to robustly and sensitively report on c-di-GMP dynamics in individual bacteria and within the physiological concentration of the signaling molecule underscores its potential value for the dissection of signal networks *in vivo*.

## CONCLUSIONS

Despite a great need and interest in following c-di-GMP and other small signaling molecules in living cells, appropriate biosensors to track c-di-GMP with high temporal resolution have been largely missing. Here, we developed two single-fluorescent protein-based biosensors, cdGreen and cdGreen2, with exceptionally large dynamic ranges of up to 70-fold. cdGreen is very useful for robust steady-state measurements in the nanomolar range. By developing cdGreen2, we have remedied the slow dissociation rate of cdGreen, which rendered it inappropriate for the recording of dynamic c-di-GMP changes *in vivo* on short time scales. Compromising on the dynamic range only marginally, we achieved rapid off kinetics, such that c-di-GMP fluctuations can be studied on physiologically relevant time scales. Both biosensors offer ratiometric readouts, providing a simple means for highly accurate measurements of ligand concentrations in individual cells *in vivo* by correcting for stochastic fluctuations in biosensor concentration. Importantly, the spectral properties of the sensors are compatible with most standard microscopy and flow cytometry setups, providing for ready adaptation and usage in most laboratories without the need to purchase specialized equipment. To demonstrate the power and versatility of this novel-generation biosensors, we have dissected the c-di-GMP-mediated genetic program that drives the *C. crescentus* bimodal life cycle and have visualized the behavior of individual cells of *Pseudomonas* during their short-term adaptation to surfaces.

While our study exclusively addresses temporal dynamics and gradients of c-di-GMP, it has been postulated that c-di-GMP operates both on a global and on a local level ^3,10,11^. Indeed, a number of studies have now shown that particular diguanylate cyclases trigger specific c-di-GMP-dependent processes through a direct interaction with their cognate effector proteins without affecting global c-di-GMP levels ^66–68^. However, insulated, subcellular c-di-GMP microenvironments have so far not been visualized. Because this will require highly sensitive or highly dynamic sensors, our tools could serve as an entry point for the development of next-generation biosensors that are able to read out the spatial organization c-di-GMP in bacteria. More generally, the design principles and functional enrichment strategies via iterative FACS developed in this study will serve as a blueprint for the development of robust genetic probes for the quantitative, dynamic *in vivo* imaging of other signaling molecules, metabolites or antibiotics. In particular, we envisage that by multiplexing various such biosensors one can investigate the interplay of different regulatory or metabolic networks and scrutinize bacterial physiology during growth in multicellular communities or during host colonization with unprecedented resolution and integrative power.

## MATERIALS AND METHODS

### Media and growth conditions

*Escherichia coli* DH5α, DH10B, TOP10 or commercial electro-competent cells (MegaX DH10b T1R and ElectroMax DH5α, see below) were used for cloning and were routinely cultivated in LB-Miller at 37 °C. The *E. coli* MG1655 derivative AKS494 (see below) was used for all biosensor experiments in *E. coli. Pseudomonas* strains were grown in LB-Lennox at 37°C overnight for pre-cultures; for cultures used for microscopy, overnight pre-cultures were diluted back 200-fold in LB-Lennox and grown at 30°C into exponential phase. *Caulobacter crescentus* strains were grown in PYE (0.2% [w/v] bacto peptone, 0.1% [w/v] yeast extract, 0.8⍰mM MgSO_4_, 0.5⍰mM CaCl_2_) at 30°C overnight for precultures; for microscopy, cultures were diluted back 50-fold in the same medium and grown for 4-6h into exponential phase at 30°C. If not otherwise mentioned, PYE contained tetracycline and was supplemented with 100 μM cumate for biosensor expression. For FtsZ depletion experiments, cultures were frown without cumate and the inducer was only included in agarose pads. Standard plates contained 1.5% [w/v] agar. For microscopy time-lapse experiments, strains were grown on 1% agarose pads with appropriate medium at 30°C. When appropriate, media were supplemented with antibiotics at the following concentrations unless stated otherwise (liquid/solid media for *C. crescentus*; liquid/solid media for *E. coli*; liquid/solid media for *P. aeruginosa*; in μg⍰ml^−1^): kanamycin (Km) (-/-; 50/50; -/-), oxytetracycline (Tc) (1/5; 12.5/12.5; -/-), chloramphenicol (-/-; 20/30; -/-), gentamycin (-/-; 20/20; 30/30), ampicillin (−/−; 100/100; -/-), carbenicillin (-/-; 100/100; -/-). The isopropy-β-D-thiogalactopyranoside (IPTG) stock was prepared in ddH_2_O at a concentration of 1⍰M. The 4-isopropylbenzoic acid (cumate) stock (100⍰mM) was prepared in 100% ethanol and used at a final concentration of 100 μM. Anhydrotetracycline (aTc) stocks (1 mM) were prepared in DMSO and used at a final concentration of 200 nM.

### Strain construction

AB607 was generated by P1-mediated transduction of the kanamycin resistance-marked *yhjH* deletion from the respective Keio collection strain ^69,70^ in strain MG1655, followed by FLP-mediated excision of the resistance cassette using pCP20 as described previously ^71^. AKS486 is a AB607 derivative carrying plasmid pAR99. AKS494 was generated by P1-mediated transduction of a kanamycin resistance-linked *pdeL’-’lacZ* translational fusion in AKS486 as described previously ^36^. A *C. crescentus* Δ*pdeA* Δ*pleD::nptII* strain was constructed by electroporation of plasmid pNPTStet-pleDnptII in UJ4454 ^44^, selection on PYE/Tc/Km plates, growth of a single colony overnight in liquid PYE/Km and plating on PYE/Km plates supplemented with 0.3% sucrose. Loss of the plasmid backbone was verified by testing for tetracycline sensitivity. A ϕCR30 lysate prepared on this strain was used to move the marked *pleD* null allele in the NA1000 *shkA*^*D369N,DD*^ *tacA*^*DD*^ background (strain AKS217)^19^ by generalized transduction. All other strains have been described previously.

### Plasmid construction

Polymerase chain reaction (PCR) was performed with Phusion DNA polymerase (NEB) in a total volume of 50 µl in GC buffer containing 10% (v/v) DMSO, 400 µM of dNTPs, 400 nM of each forward and reverse primer, 100-200 ng of DNA template and 0.4 µl of Phusion DNA polymerase. pAR81 was constructed by PCR-amplification of *yhjH* from an E. coli MG1655 colony with primers 7323/7324 and cloning in pNDM220 ^72^ via BamHI/XhoI. pAR94 was constructed by PCR-amplifying a *parMR*-containing fragment from pNDM220 using primers 6801/6803, a *cat*-containing fragment from pKD3 ^71^ with primers 6802/6804, fusion of the two fragments by SOE-PCR using primers 6801/6804 and cloning in pNDM220 via PscI/AatII. pAR99 was constructed by subcloning a fragment carrying *yhjH* from pAR81 in pAR94 via BamHI/XhoI. pATTPtetycgR1 was constructed as follows. *ycgR* was PCR-amplified with primers 12056/12057 using an *E. coli* TOP10 colony as template. *Ptet-tetR* was PCR-amplified with primers 11469/12059 using plasmid pAR83 as template, followed by a second PCR with primers 12058/12059 using the first PCR product as template. pAR83 was a gift from Attila Becskei (Biozentrum, University of Basel) and encodes an autoregulatory P*tet*-TetR system. The two fragments, *ycgR* and *Ptet-tetR*, were joined by SOE-PCR, digested with PciI/AatII and cloned in pATT-Dest ^34^ digested with the same enzymes. pATT-Dest was a gift from David Savage (Addgene plasmid #79770). pATTPtetycgR2, which contains a unique EcoRI site, was constructed by PCR-amplification of a fragment from pATTPtetycgR1 using primers 12157/12059, digestion with PciI/XbaI and cloning in the same sites of pATTPtetycgR1. pATTPtetycgR3, which contains a strong ribosome binding site upstream of *ycgR*, was constructed by PCR-amplification of a fragment from pATTPtetycgR2 using primers 12435/12436, digestion with AfeI/EcoRI and cloning in the same sites of pATTPtetycgR2. pBldDtemp was generated by PCR-amplification of *bldD* with primer pairs 14252/14253 and 14254/14255 from plasmid pIJ10663 ^35^, fusion of the two fragments by SOE-PCR lacking primers and cloning in EcoRI/SpeI-digested pATTPtetycgR3 using Gibson assembly ^73^. pBldDtemp contains a tandem fusion of *bldD* with BsaI restriction sites in between *bldD* copies to allow domain/linker insertion using Golden Gate cloning ^74^. p2H12 was isolated as a 4th generation c-di-GMP sensor by FACS (see below). p2H12Matry was constructed by PCR-amplification of mScarlet-I from pBAD24_VCA0107_mScarlet-I_Cterm_04 (a gift from Johannes Schneider, group of Marek Basler, Biozentrum, University of Basel) with oligos 15688/15689 and cloning in KpnI-digested p2H12 via Gibson assembly. p2H12Matry-blind was constructed in two steps. First two fragments were PCR-amplified with primer pairs 15631/15633 and 15632/15634 from plasmid p2H12 and cloned in KpnI/EcoRI-digested p2H12 using Gibson assembly. Then, the plasmid was digested with KpnI and mScarlet-I was PCR-amplified from p2H12Matry with oligos 15688/15689 and assembled via Gibson cloning. p2H12ref was constructed by PCR-amplification of mScarlet-I from pBAD24_VCA0107_mScarlet-I_Cterm_04 with oligos 15549/15550 and cloning in SpeI-digested p2H12 via Gibson assembly. For construction of pConRef-2H12, oligos 16066/16067 were phosphorylated and annealed, and ligated to p2H12ref digested with XhoI/NdeI, replacing the original *P*_*tet*_ promoter and most of *tetR*. pConRef-2H12.D11 was constructed by subcloning a fragment encoding 2H12.D11 from pQFmcs-2H12.D11 (see below) in pConRef-2H12 via EcoRI/SpeI. pQFmcs was constructed by annealing phosphorylated oligos 14670 and 14671 and cloning the resulting dsDNA fragment in pQF ^75^ digested with EcoRI/SpeI. pQFmcs-2H12 was constructed by subcloning a fragment encoding 2H12 from p2H12 in pQFmcs via EcoRI/SpeI. p2H12.D11 and pQFmcs-2H12.D11 were obtained as described below. pQFmcs-2H12.D11-scarREF was constructed by subcloning a EcoRI/NheI fragment from pConRef-2H12.D11 in pQFmcs-2H12.D11 digested with EcoRI/SpeI. pAK206-2H12.D11 was constructed by subcloning a EcoRI/SpeI-fragment encoding 2H12.D11 from pQFmcs-2H12.D11 in pAK206 ^76^ digested with NheI/EcoRI. Final pAK206-2H12.D11 showed an unusual SpeI/NheI scar (ACTAGTAGC, instead of the expected ACTAGC), leaving the SpeI site intact. For construction of pBBR15-2H12.D11, oligos 16583/16584 were used for round-the-horn PCR on pAK206-2H12.D11 as a template. pBBR15-2H12.D11_v2 (which has a second SpeI on the backbone eliminated) was constructed by PCR-amplification of pBBR15-2H12.D11 with oligos 18655/18656 and circularization of the PCR product using Gibson assembly. pBBR15.2-2H12.D11opt was constructed by cloning a high-GC-content-optimized DNA fragment encoding a variant of 2H12.D11 (synthesized by Twist Bioscience) in pBBR15-2H12.D11_v2 digested with NheI/SpeI via Gibson assembly. pBBR15.2-2H12.D11opt-scar was constructed by subcloning a SpeI/NheI-fragment encoding mScarlet-I from pConRef-2H12 in pBBR15.2-2H12.D11opt digested with SpeI and dephosphorylated. pET28-2H12, pET28-2H12-Matry and pET28-2H12.D11 were constructed by PCR-amplification of a fragment from p2H12, p2H12-Matry or p2H12.D11, respectively, using primer pair 15629/15630 and cloning in pET28a via NdeI/SacI. pNPTStet-pleDnptII was constructed by PCR amplification of *nptII* conferring kanamycin resistance from pAK405 ^77^ using primer pair 18357/18358, two fragments encoding *pleD* up- and downstream regions using primer pairs 18403/18404 and 18405/18406 from *C. crescentus* genomic DNA, respectively, and cloning of the three fragments in pNPTStet ^19^ digested with SpeI/SphI using Gibson assembly.

### Biosensor libraries constructions: cdGreen

For the initial library (“library 1”), cpGFP was PCR-amplified from pTKEI-Tre-C04 ^34^ (Addgene plasmid #79754, a gift from David Savage) using standard PCR conditions with the reaction mix containing 67 nM each of the following primers: 14292, 14293, 14294, 14295, 14296, 14297, 14298, 14299, 14300, 14301, 14302, 14303. The following cycling program was used: 98 °C for 2 min; 98°C for 10 s, 55°C for 10 s, and 72°C for 30 s (30 cycles); 72 °C for 1 min; and hold at 10 °C. After amplification, 1 ul of DpnI was added to the reaction and incubated at 37°C for 1h to digest the DNA template, followed by purification of the PCR product using a NucleoSpin Gel and PCR Clean-Up Kit (Macherey–Nagel) with elution in 15 µl NE elution buffer. The library was constructed using Golden Gate Assembly in a total volume of 50 µl in T4 DNA ligase buffer containing 250 ng of plasmid pBldDtemp, 1.75 µg of purified cpGFP PCR product, 1000 units T4 DNA ligase (NEB) and 30 units BsaI-HFv2 (NEB). The reaction was incubated 90 min at 37°C, 15 min at 55°C and 20 min at 80°C and purified using a NucleoSpin Gel and PCR Clean-Up Kit (Macherey– Nagel) with elution in twice 15 µl ddH_2_O. The plasmid library was concentrated in a Eppendorf Concentrator plus (20 min, 30°C, V-AQ) to approximately 5 µl and 4 µl were transformed into 100 µl of ElectroMax DH5α-E Competent Cells (Cat# 11319019, Thermo Fisher Scientific). After 30 min of recovery at 37°C in 4 ml of SOC medium, cells were added to 400 ml LB containing 100 µg/ml carbenicillin. 10 µl were plated on LB plates containing 100 µg/ml ampicillin to calculate the library size and the remainder was incubated at 37°C with shaking (180 rpm) overnight. The library was estimated to comprise 1.9 10^6^ individual clones. Plasmid library DNA was purified from twice 5 ml of culture using a GenElute Plasmid Miniprep Kit (Sigma-Aldrich) with elution in twice 20 µl of ddH_2_O, followed by concentration in an Eppendorf Concentrator plus (20 min, 30°C, V-AQ). The library was transformed in strain AKS494 by electroporation using a BioRad GenPulser (1.75kV, 25µFD, 400 Ohms) and transformations (estimated to contain >2 10^8^ clones) were outgrown in 400 ml LB containing 100 µg/ml carbenicillin, 6 µg/ml chloramphenicol and 200 nM aTc. Libraries for linker optimization (libraries 2, 3 and 4) were constructed similarly, except that different primer sets and different templates were used for PCR amplification of cpGFP, GGA was performed by incubation of the reactions overnight at 37°C and MegaX DH10B T1R Electrocomp Cells (Cat# C640003, Thermo Fisher Scientific) were used for cloning. In each round of linker optimization, two amino acid changes at a time (one in the N-terminal and one in the C-terminal linker each) were allowed in all possible combinations. The best-performing biosensors were selected for the next linker optimization round, the “optimized” amino acids were retained and the other positions were randomized two at a time as decribed above. Library 2: oligos 14812, 14813, 14814, 14815, 14816, 14817, 14818, 14819, 14820, 14821. 14822, 14823, 14824 and 14825; pBldD2 as template. Library 3: oligos 14990, 14991, 14992, 14993, 14994, 14995, 14996, 14997, 14998, 14999, 15000 and 15001; pBldDX19-13 as template. Library 4: oligos 15202, 15206, 15207, 15208 and 15209; pBldD-A8 as template. Libraries 2-4 all contained > 10^6^ individual clones and were passaged through ElectroMax DH5α-E Competent Cells (for EcoKI methylation) before transformation in the final screening strain AKS494 as described above. All libraries transformed in AKS494 were outgrown overnight and used to inoculate cultures for FACS the next day.

### Fluorescence-activated cell sorting (FACS)

Overnight cultures were diluted 100-fold in 5 ml of LB supplemented with 100 µg/ml carbenicillin, 6 µg/ml chloramphenicol and 200 nM aTc with (1 mM) or without IPTG and grown for 2h at 37°C in a drum roller (125 rpm). 500 µl of culture were added to 5 ml of PBS and kept on ice. All sorts were performed using a BD FACSAria III cell sorter equipped with a 488-nm blue laser, a 495 nm LP mirror and a 514/30 nm BP filter for detection of GFP fluorescence at the FACS Core Facility, Biozentrum, University of Basel. Cells were sorted in “purity” mode in 5 ml of LB containing 100 µg/ml carbenicillin, 6 µg/ml chloramphenicol and 200 nM aTc. For initial naïve libraries, between 10^6^ and 10^7^ cells grown without IPTG showing a positive GFP signal were sorted and grown overnight at 37°C in a roller drum (125 rpm). To set a threshold for GFP-positive cells, AKS494 harboring plasmid pBldDtemp was used as a negative control. After these initial sorts, one “selection cycle” consisted of one round of FACS of cultures grown without IPTG (high c-di-GMP levels) and sorting for high fluorescence, followed by outgrowth of sorted cells, back-dilution, growth in medium with 1 mM IPTG (low c-di-GMP levels) and sorting for high fluorescence. The gates for high and low fluorescence were defined as containing the top or bottom, respectively, 1-5% of the main GFP distribution. In case there was a clear GFP-negative subpopulation (probably representing cells that have lost the biosensor plasmid entirely), both this small subpopulation and the lowest 1-5% of the main population were sorted in the “c-di-GMP low” step. After 6-10 selection cycles, depending on the specific library, single clones were isolated after an additional FACS round for “high c-di-GMP” and individually tested for their response to the high and low c-di-GMP regimes as described above for libraries. For clones showing the desirable c-di-GMP-dependent response in GFP fluorescence intensity, plasmids were isolated using a GenElute Plasmid Miniprep Kit (Sigma-Aldrich) and sequenced using standard primers EGFP-C-for (GTCCTGCTGGAGTTCGTG) and EGFP-N-rev (GCTTGCCGTAGGTGGCATC) at Microsynth (Balgach, Switzerland).

### Biosensor library construction and FACS: 2H12-Aff2 and 2H12-Aff8

A library of variants with partially randomized cdG-binding motifs in the 5’-encoded bldDCTD portion of plasmid p2H12 were generated by PCR amplification of two fragments from p2H12 using primer pairs 15937/15939 and 15634/15938, joining of the two fragment via SOE-PCR using primers 15634 and 15939 and cloning in p2H12 digested with EcoRI/KpnI via Gibson assembly. The Gibson assembly reaction was purified using a NucleoSpin Gel and PCR Clean-Up Kit (Macherey–Nagel) with elution in twice 15 µl ddH_2_O and the entire reaction was transformed into 100 µl of ElectroMax DH5α-E Competent Cells (Cat# 11319019, Thermo Fisher Scientific) as described above. The library comprised ca. 500’000 clones and was subsequently transformed in AKS494 as described above and subjected to FACS. Iterative FACS to enrich for functional sensors was performed essentially as described above for cdGreen with the critical exception that the “c-di-GMP low” condition was achieved by adding 20 µM IPTG instead of 1 mM IPTG. This IPTG regime was chosen to enrich for cdGreen variants that would “turn off” at intermediate c-di-GMP concentrations, i.e. at higher c-di-GMP concentrations compared to the original biosensor, with the expectation that such variants would display a lower K_d_, possibly due to faster off kinetics. After 5 selection cycles, single colonies were isolated and tested individually. Two plasmids, p2H12-Aff2 and p2H12-Aff8, were isolated and sequenced and chosen for further studies.

### Combinatorial screening of c-di-GMP motif variants in *C. crescentus*: cdGreen2

Individual mutations in motif 1 (SQRGD) or motif 2 (RQDD) in the BldD c-di-GMP-binding site were then constructed in either the 5’ or 3’ *bldDCTD* part of cdGreen. For mutations in the 5’ part, SOE-PCRs were performed with flanking primers 15634 and 15939 and mutagenic primer pairs 16130/16131, 16132/16133, 16134/16135 or 16136/16137 and p2H12 as template and cloning in p2H12 via EcoRI/KpnI and Gibson assembly. For mutations in the 3’ *bldDCTD*, SOE-PCRs were performed with flanking primers 15138 and 15139 and mutagenic primer pairs 16130/16131, 16132/16133, 16134/16135 or 16136/16137 and p2H12 as template and cloning in p2H12 via KpnI/SpeI and Gibson assembly. To introduce both motif 1 and motif 2 mutation of 2H12-Aff2 or 2H12-Aff8 in the 3’ bldDCTD, SOE-PCRs with 15138 and 15139 and mutagenic primer pairs 16188/16190 or 16189/16191, respectively, and p2H12 as template and cloning in p2H12 via KpnI/SpeI via Gibson assembly. Mutations in the 5’- and 3’-encoded *bldDCTD*s were combined by subcloning 3’-encoded *bldDCTD* variants in p2H12 derivative encoding 5’-encoded *bldDCTD* variants via KpnI/SpeI. cdGreen variants were subcloned into pQFmcs-2H12 (replacing original cdGreen) via EcoRI/SpeI. Individual plasmids were then transformed in *C. crescentus* NA1000, resulting strains were grown overnight in 5 ml PYE supplemented with 1 ug/ ml oxytetracycline and 100 uM cumate, diluted 20-fold in the same medium and grown for another 4-6 h before being imaged on 1% agarose PYE pads by microscopy using a DeltaVision system. Images were visually inspected for late predivisional cells that showed asymmetric distributions of GFP fluorescence in the two (future) daughter cell compartments. A single cdGreen derivative that harbors the RPAD sequence in c-di-GMP-binding motif 2 in both BldD_CTD_ protomers, named cdGreen2, met this criterion and was chosen for further characterization.

### Expression and purification of biosensors cdGreen, cdGreen-Matry and cdGreen2

cdGreen, cdGreen-Matry and cdGreen2 were overexpressed in *Escherichia coli* BL21(DE3). Cells were grown at 37°C in terrific broth medium containing 50⍰μg⍰ml^-1^ kanamycin to an OD_600_ ⍰of 0.8–1.0. Protein expression was induced by 0.1⍰mM IPTG, then the temperature was reduced to 20°C and cells were harvested after further overnight cultivation. After lysis of the cells with a sonicator in 50 mM Tris-HCl pH 8.0, 250 mM NaCl, 40 mm imidazole, the soluble biosensor was purified by immobilized metal-affinity chromatography on a Ni Sepharose column (Cytiva) and then subjected to size exclusion chromatography on a Superdex S200 column (Cytiva) in sensor buffer (25 mM Tris-HCl pH 7.5, 150 mM NaCl, 10 mM MgCl2, 10 mM KCl, 5 mM 2-Mercaptoethanol). The protein-containing fractions were pooled and concentrated in Amicon Ultra units (Merck). Protein yields were about 20 mg L^-1^.

### Isothermal titration calorimetry (ITC)

ITC experiments were performed at 30°C in sensor buffer, using a Microcal iTC200 instrument. The concentration of cdGreen in the calorimeter cell was 25 µM and the concentration of cyclic-di-GMP in the injector syringe was 1000 µM. 1 × 0.4 µl and 19 × 1.8 µl injections of cyclic-di-GMP were performed with stirring at 750 rpm, and the resulting heats of injection were monitored with an inter-injection spacing of 180 s, and an averaging time of 1 s. As a control or the heat of dilution and dissociation of cyclic-di-GMP, we performed an experiment that was identical in all respects apart from the omission of cdGreen from the solution in the calorimeter cell. The raw data were baseline-corrected, integrated, and analysed using the web version of the software Affinimeter ^78^. To correct for the heat of dilution and dissociation of cyclic-di-GMP, the integrated heats of injection for cyclic-di-GMP injected into buffer were subtracted from the integrated heats of injection for the injection of cyclic-di-GMP into cdGreen. The resulting heats of injection were fitted to the model described below, specified using the model-builder function of Affinimeter, with parameter errors determined automatically in the software by the Jackknife method.

### ITC: Model fitting

We expected four binding sites for cyclic-di-GMP. The general scheme for the binding equilibrium is therefore the stepwise association of four cyclic-di-GMP molecules (cdG) to the protein (P):

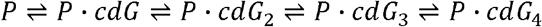

Macroscopic association constants and enthalpy change for binding can be assigned to each step (*K*_*1*_ – *K*_*4*_, Δ*H*_*1*_ – Δ*H*_*4*_, for each step left to right in the equilibrium above). The ITC titration of cyclic-di-GMP shows two clear phases, one at a molar ratio of 2 cyclic-di-GMP : 1 protein, and a second at a molar ratio of 4 cyclic-di-GMP : 1 protein, with different enthalpy changes for binding evidenced by the different heat of injection in the initial plateau of each phase (ca -16 kcal/mol for the first phase, -7 kcal/mol for the second phase). This confirms the anticipated total stoichiometry of 4 cyclic-di-GMP : 1 protein. The simplest equilibrium scheme that can fit such data comprises two sets of two equivalent binding sites. In terms of macroscopic association constants, this model fits *K*_*1*_ and *K*_*3*_, and sets *K*_*2*_ = 0.25^*^ *K*_*1*_ and *K*_*4*_ = 0.25^*^ *K*_*3*_ according to the expected statistical factor for two equivalent binding sites. The microscopic binding constants (site constants) for each set of equivalent sites can be obtained from *K*_*micro1,2*_ = 0.5 *K*_*1*_ and *K*_*micro3,4*_ = 0.5 *K*_*3*_. In terms of enthalpy changes for binding, this model fits Δ*H*_*1*_ and Δ*H*_*3*_, and sets Δ*H*_*2*_ = Δ*H*_*1*_ and Δ*H*_*4*_ = Δ*H*_*3*_. The model is not a perfect fit to the data. It is probable that the underlying equilibrium is more complicated (e.g. having non-equivalent binding sites, and cooperativity) but inclusion of further fitting parameters without additional information or data would result in an over-parameterised fit. The fit gave *K*_1_ = 3.98 × 10^8^ M^-1^ (± 0.64 × 10^8^ M^-1^), *K*_3_ = 6.05 × 10^5^ M^-1^ (± 1.18 × 10^5^ M^-1^), Δ*H*_*1*_ = -16.2 kcal/mol (± 0.1 kcal/mol), and Δ*H*_*3*_ = -7.5 kcal/mol (± 0.3 kcal/mol). A concentration correction factor of 0.96 was fitted for the protein. These macroscopic association constants correspond to microscopic *K*_*D*_ (site constant) values of 5.0 nM for the higher-affinity set of two sites, and 3.3 µM for the lower-affinity set of two sites.

### *In vitro* c-di-GMP biosensor characterization

All measurements were taken in a SynergyH4 plate reader (BioTek) with protein and c-di-GMP in a total volume of 200 µl sensor buffer using 96-well black/clear flat bottom plates (Costar) at 30°C. Emission and excitation spectra of purified cdGreen, cdGreen-Matry and cdGreen2 shown in Figures 2a and 3d were recorded at 30°C at a protein concentration of 1 µM with excitation and emission wavelengths detailed in the figures or figure legends. C-di-GMP was added to 30 µM final concentration from a 1 mM stock solution in ddH_2_O and all reactions were let to equilibrate at 30°C for 2h before taking measurements. Scans were at 1 nm resolution using the minimal bandwidth of 9 nm. For scans shown in Figures 2b and 3e, and dose-response curves shown in Figures 2c and 3f, protein concentration was 150 nM and c-di-GMP concentrations as indicated in figures or figure legends. Sensor and c-di-GMP were mixed and let equilibrate for 7h at 30°C before measurements were taken, which was sufficient to reach binding equilibrium for all c-di-GMP concentrations tested (Supplementary Fig. 6). The multiple data points shown for each c-di-GMP concentration are repeated measurements of the same well every minute over a 10-min period. Excitation and emission wavelengths are specified in the figures or figure legends (minimal bandwidth of 9 nm). K_d_s were fitted using GraphPad Prism 9 software and the “[Inhibitor] vs. response -- Variable slope (four parameters)” model. Other nucleotides tested for cdGreen were purchased from Jena Bioscience and tested at a concentration of 10 µM with 150 nM sensor with (10 µM) or without c-di-GMP after 5 h of equilibration at 30°C. For stoichiometric titrations experiments, protein concentration was 25 µM and c-di-GMP was added from a 5 mM stock solution in sensor buffer to give the final concentrations indicated in the figures or figure legends. Sensor and c-di-GMP were mixed and let equilibrate for 30 min at 30°C before measurements were taken. For “dissociation by dilution” experiments to determine k_off_ rate constants, 7.5 µM of sensor was mixed with 200 nM (for cdGreen) or 300 nM (for cdGreen2) c-di-GMP and let equilibrate for 7h at 30°C. Solutions were then diluted 50-fold in sensor buffer (4 µl in 200 µl sensor buffer; final c-di-GMP concentration of 4 nM for cdGreen and 6 nM for cdGreen2) and measurements were started immediately (ca. 10 sec dead time). For cdGreen, a control was included in parallel, where cdGreen at 7.5 µM without added c-di-GMP was diluted in the same way as the “plus c-di-GMP” sample in sensor buffer, and the baseline response of the control was subtracted from the “plus c-di-GMP” response. This correction was done because dilution of cdGreen showed an initial increase of fluorescence intensity after dilution, which was largely attributable to a protein dilution effect. cdGreen2 did not show such a behaviour, so it was not necessary to include such a control for this biosensor. k_off_ was fitted using GraphPad Prism 9 software and the “Dissociation - One phase exponential decay” model. Global estimation of k_on_, k_off_ and K_d_ was performed using association kinetics with three ligands concentrations (25 μM, 12.5 μM and 6.25 μM) at a cdGreen or cdGreen2 concentration of 150 nM. 5 μl of c-di-GMP were added from 40-fold stock solutions to 195 μl cdGreen or cdGreen2 and measurements were started immediately (dead time of ca. 10 sec). Association and dissociation rate constants and the equilibrium dissociation rate constant were fitted globally using the “Association kinetics (two ligand concentration)” model in GraphPad Prism 9.

### Flow cytometry

Exponentially growing cultures were sampled in ddH_2_O and analyzed on a BD LSR Fortessa using Diva software (BD Biosciences). After standard SSC-H/FSC-H and SSC-H/SSC-W gating for singlets, cells were gated for mScarlet-I-positive cells using a yellow-green 561 nm laser, a 600 nm long-pass mirror and a 610/20 nm band-pass filter, and cells were analyzed for their emission signals (long-pass mirror: 505 nm; band-pass filter: 512/25 nm) following excitation with the violet 405 nm or the blue 488 nm laser. 50’000 events in the mScarlet-I-positive gate per sample were collected. Data were analyzed using FlowJo v10.0.6 (FlowJo LLC). Ratios of emission signals upon excitation with 488 and 405 nm were calculated and plotted in FlowJo using the built-in “Derived parameters” tool.

### Microscopy

Images were acquired using a DeltaVision system with softWoRx 6.0 (GE Healthcare) on a Olympus IX71 microscope equipped with a pco.edge sCMOS camera and an UPlan FL N 100X/1.30 oil objective (Olympus). Signals for the cdG sensor were acquired using 390/18 nm (DAPI) and 475/28 nm (FITC) excitation filters and a 523/36 nm (FITC) emission filter. mScarlet was imaged using 575/25 nm excitation and 632/60 nm emission (A594) filters. For time lapse experiments, the following illumination settings were applied (Excitation/Emission, Exposure time, Power, Imaging interval) for experiments shown in Figure 3c: FITC/FITC, 10 ms, 50%, 5 min; DAPI/FITC, 5 ms, 50%, 5 min. For Figure 3h and Supplementary Movie 1: FITC/FITC, 100 ms, 2%, 20 s; A594/A594, 100 ms, 2%, 20 s. For Figure 4: FITC/FITC, 15 ms, 50%, 5 min; A594/A594, 25 ms, 50%, 5 min. For Figure 5: FITC/FITC, 150 ms, 10%, 5 min; DAPI/FITC, 100 ms, 5%, 5 min; A594/A594, 150 ms, 5%, 5 min. For Supplementary Figure 3: FITC/FITC, 15 ms, 50%, 5 min; DAPI/FITC, 10 ms, 50%, 5 min. For Supplementary Figure 4c: FITC/FITC, 10 ms, 50%, 5 min; DAPI/FITC, 5 ms, 50%, 5 min; A594/A594, 100 ms, 100%, 5 min. For Supplementary Figure 5a: FITC/FITC, 150 ms, 5%, 5 min; DAPI/FITC, 100 ms, 5%, 5 min. Data shown in Figure 2e was acquired using NIS Elements on a Nikon Eclipse Ti2 inverted microscope equipped with a Hamamatsu ORCA-Flash4.0 V3 Digital CMOS camera and a 100X/1.45 Nikon Plan Apo Lambda 100x Oil Ph3 DM objective. Signals for the cdG sensor were acquired using 395/25 nm and 470/24 nm excitation filters and a 515/30 nm emission filter. mScarlet was imaged using 575/25 nm excitation and 632/30 nm emission filters. Cells were imaged within 15 min after spotting on agarose pads. Images were analyzed using the Fiji ^79^ plugin MicrobeJ ^80^.

### Time-lapse image analysis

Time-lapse movies were visually inspected using Fiji ^79^ and trimmed to exclude later frames were cells overlapped. Subsequently, cells were segmented and tracked using the DeLTA 2.0 ^81^ deep-learning based pipeline. For *P. aeruginosa* we used the default pre-trained model for both segmentation and tracking. For *C. crescentus* we retrained the segmentation model using our own training data (the default tracking model was used). Training data was obtained by segmenting cells using the Ilastik pixel classification pipeline ^82^, followed by postprocessing using custom written Python code, and manually curation using Napari (https://zenodo.org/record/3555620). Further data analysis was performed using custom written Python code (available here: https://github.com/simonvanvliet/cdg_dynamics_caulobacter). Visual inspection of the processed data showed that segmentation errors were extremely rare, but tracking errors occurred regularly, especially at later time points. We therefore implemented and automated screening method to filter out erroneous cell tracks. Specifically, we only included pairs of sister cells which were both tracked for at least 12 frames (1h) for *C. crescentus* or 8 frames (40 min) for *P. aeruginosa*. We only consider frames where data for both sister cells was available (i.e., the track of the longest-lived sister was trimmed to the same length of the shortest-lived sister). Moreover, we excluded pairs were one or both cells showed unexpectedly large jumps in cell length between two frames (a decrease of more than 8% or an increase of more than 8% (*C. crescentus*) or 12% (*P. aeruginosa*)). Finally, we excluded cell pairs where the summed length of the sister cells at birth deviated strongly (a decrease of more than 6% or an increase of more than 20%) from the cell length of the mother cell just before division. For the remaining pairs of cells, the cdG level was estimated as the mean fluorescent intensity in the GFP/FITC channel. We then grouped sister cells into two classes: the cdG-high and cdG-low class, based on the average cdG level over the full lifetime of the cells.

## ACKNOWLEDGEMENTS

We thank Mark Buttner, David Savage, Attila Becskei, Johannes Schneider and Marek Basler for plasmids; Timothy Sharpe (Biophysics Core Facility, Biozentrum, University of Basel) for expertise support with ITC experiments and advice on stoichiometric titrations; Janine Bögli and Stella Stefanova (FACS Core Facility, Biozentrum, University of Basel) for their technical support with FACS; Raphael Dias Teixeira for preparation of illustrations shown in Fig. 1b,c; and Fabienne Hamburger for help with cloning and strain construction. This work was supported by the Swiss National Science Foundation grants 310030B_185372 and 310030_208107 to U.J.; and by NCCR AntiResist funded by Swiss National Science Foundation grant 51NF40_180541 to U.J. S.v.V. was funded by an Ambizione grant from the Swiss National Science Foundation (grant nr. PZ00P3_202186) and by the University of Basel.

## AUTHOR CONTRIBUTIONS

U.J. and A.Ka. conceived and designed experiments. A.Ka. performed experiments. A.Ka., S.v.V. and U.J. analyzed data. R.P.J., A.R., A.Kl. and T.M. contributed materials. U.J. and A.Ka. wrote the draft with input from all authors.

## FIGURE LEGENDS

**Supplementary Fig 1 – SDS-PAGE analysis with Coomassie staining of purified biosensors**

a. SDS-PAGE analysis with Coomassie staining of N-terminally His_6_-tagged cdGreen (lane 1) and cdGreen-Matry (lane 2) after NiNTA purification and size exclusion chromatography. A protein standard was run in lane 3 and molecular weights of relevant bands are indicated.

b. SDS-PAGE analysis with Coomassie staining of N-terminally His_6_-tagged cdGreen2 (lane 1) after NiNTA purification and size exclusion chromatography. A protein standard was run in lane 2 and molecular weights of relevant bands are indicated.

**Supplementary Figure 2 – Specificity and stoichiometry of c-di-GMP binding to biosensors**

a. Response of purified cdGreen (150 nM) to different nucleotides (10 μM) was measured, with (10 μM) or without additional c-di-GMP. Responses were recorded on a microplate reader and values are 530 nm emission ratios upon excitation with 497 nm and 405 nm, normalized to the value of this signal obtained with cdGreen with 10 μM c-di-GMP. Shown are means of N = 6 repeated measurements at equilibrium.

b. Stoichiometric titration of cdGreen. cdGreen (25 μM) was incubated with varying concentrations of c-di-GMP (see Figure) to equilibration and responses were recorded on a microplate reader using emission at 530 nm upon excitation at 497 nm as a readout. Shown are means of N = 6 repeated measurements at equilibrium.

c. Isothermal titration calorimetry (ITC) of cdGreen with c-di-GMP. For details, see Materials and Methods and the Figure.

d. Stoichiometric titration of cdGreen2 with c-di-GMP. Experiments were carried out as described for cdGreen in panel b. Shown are means of N = 6 repeated measurements at equilibrium.

e. Association kinetics of cdGreen (for details, see Materials and Methods). Values represent 530 nm emission ratios upon excitation with 497 nm and 405 nm. Shown are individual data points from N = 3 replicates. Note that GraphPad Prism 9 failed to globally fit kinetic parameters based on these data.

f. Association kinetics of cdGreen2 (for details, see Materials and Methods). Values represent 530 nm emission ratios upon excitation with 497 nm and 405 nm. Shown are individual data points from N = 3 replicates. Different c-di-GMP concentrations used are as in panel e.

**Supplementary Figure 3 – *In vivo* performance of biosensor cdGreen in *C. crescentus***

a. Time-lapse series (5-min intervals) of strain NA1000 carrying plasmid pQFmcs-2H12. Individual channels in black/white (BW) or pseudo-colors (C, M, Y) are shown as well as an overlay of the three pseudo-colored channels. The scale bar indicates 2 µm. The yellow arrow indicates the stalked pole.

b. Time-lapse series (15-min intervals) of strain NA1000 rcdG^0^ carrying plasmid pQFmcs-2H12. Individual channels in black/white (BW) or pseudo-colors (C, M, Y) are shown as well as an overlay of the three pseudo-colored channels. The scale bar indicates 2 µm.

**Supplementary Figure 4 – Screening and isolation of biosensor cdGreen2**

a. Microscopy snapshots of *C. crescentus* strains carrying different 2H12 derivatives (see Materials and Methods for details). Variants of c-di-GMP binding motifs in the N-terminal BldD CTD (1) and/or the C-terminal BldD CTD (2) are indicated in the top left corner of each panel. A single late predivisional showing asymmetric c-di-GMP levels is highlighted by an orange box (variant “1: RPAD, 2: RPAD” aka cdGreen2).

b. More examples of late predivisional cells with asymmetric c-di-GMP levels in the “1: RPAD, 2: RPAD” sample from the same screening setup shown in panel a.

c. Time-lapse series of a NA1000 strain chromosomally expressing mCherry-tagged SpmX, serving as a ST cell-specific marker, and carrying plasmid pQFmcs-2H12.D11. Time stamps are in minutes. The scale bar represents 2 µm.

d. Time-lapse series of strain NA1000 carrying plasmid pQFmcs-2H12.D11. Individual channels in black/white (BW) or pseudo-colors (C, M, Y) are shown as well as an overlay of the three pseudo-colored channels. The scale bar indicates 2 µm. Note that this time-lapse series is part of the series shown in Figure 3c.

**Supplementary Figure 5 – C-di-GMP dynamics upon FtsZ depletion in *C. crescentus***

a. Time-lapse series of a NA1000 derivative carrying chromosomally encoded *dcas9* under control of the cumate-inducible promoter P_Q5_, a plasmid constitutively expressing a sgRNA targeting *ftsZ*, and plasmid pQFmcs-2H12.D11 expressing cdGreen2. Images are overlays of the cdGreen2 FITC/FITC signal pseudo-colored in yellow and the DAPI/FITC signal pseudo-colored in magenta. Time stamps indicate minutes after spotting on agarose pads. Note that FtsZ depletion was initiated by spotting on agarose pads containing cumate. Contrast and brightness settings are the same for all images.

b. Same as in panel a, showing an example, in which an elongated cell manages to divide late after triggering FtsZ depletion. Contrast and brightness settings are the same for all images, except for the first two time points, which were manually adjusted to better see the biosensor signal (note that expression of cdGreen was only induced upon spotting on agarose pads). The arrows point at the septum/division site.

c. Flow cytometry analysis of indicated strains. The rcdG^0^ *Plac-ydeH* strain lacks all endogenous diguanylate cyclases and phosphodiesterases, but harbors a chromosomal, IPTG-inducible copy of the heterologous diguanylate cyclase DgcZ from *E. coli*; no IPTG (-), 1 mM IPTG (+). For each event measured, the 488 nm/ 405 nm ratio is given. On top, the gates used for classification as cells with low or high c-di-GMP (see panel d) are indicated.

d. Quantification of fractions of cells with low c-di-GMP levels (as defined by the gates shown in panel c) for indicated strains. Shown are means and standard deviations of three biological replicates per strain and condition.

**Supplementary Figure 6 – Equilibration of c-di-GMP/biosensor complexes for *in vitro* experiments**

Indicated concentrations of c-di-GMP were added to 150 nM biosensor and the biosensor signals (excitation at 497 nm, emission at 530 nm) were recorded over time at 30°C.

**Supplementary Movie 1 – High-temporal-resolution imaging of c-di-GMP levels in *C. crescentus***

Time-lapse movie of *C. crescentus* cells carrying pQFmcs-2H12.D11-scarREF imaged in 20-sec intervals over ∼3 hours at 40 frames per seconds. The right panel shows the phase contrast channel; the left panel shows an overlay of the cdGreen2 FITC/FITC signal pseudo-colored in yellow, the mScarlet-I signal pseudo-colored in magenta and the phase contrast signal pseudo-colored in cyan.

