## Supplementary figures and images for "A Novel Biosensor Reveals Dynamic Changes of C-di-GMP in Differentiating Cells with Ultra-High Temporal Resolution"

### Supplementary Figure 1

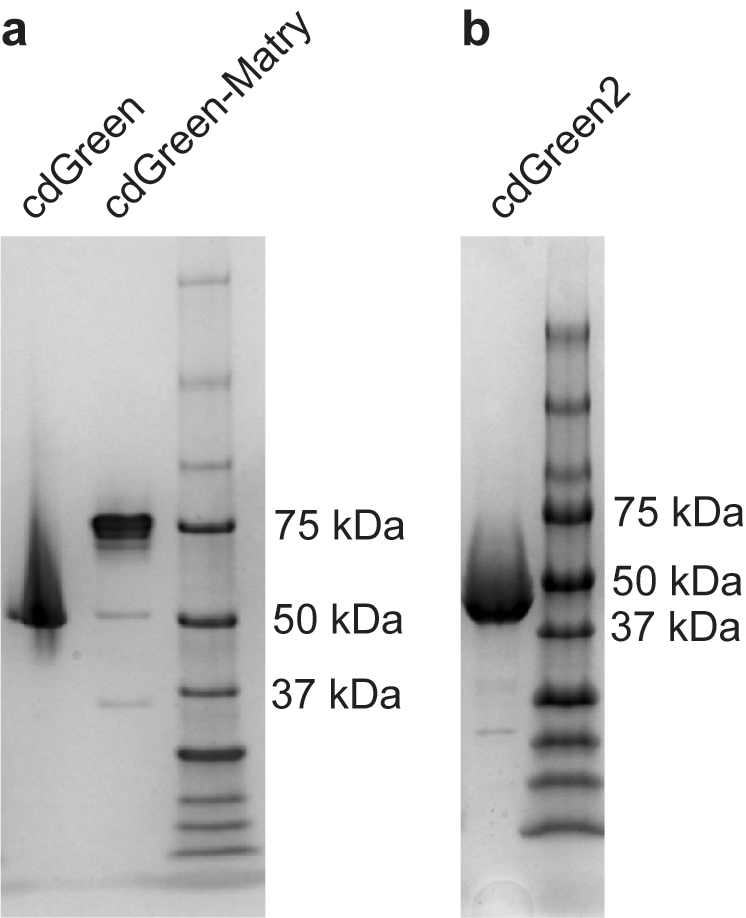

### Supplementary Figure 2

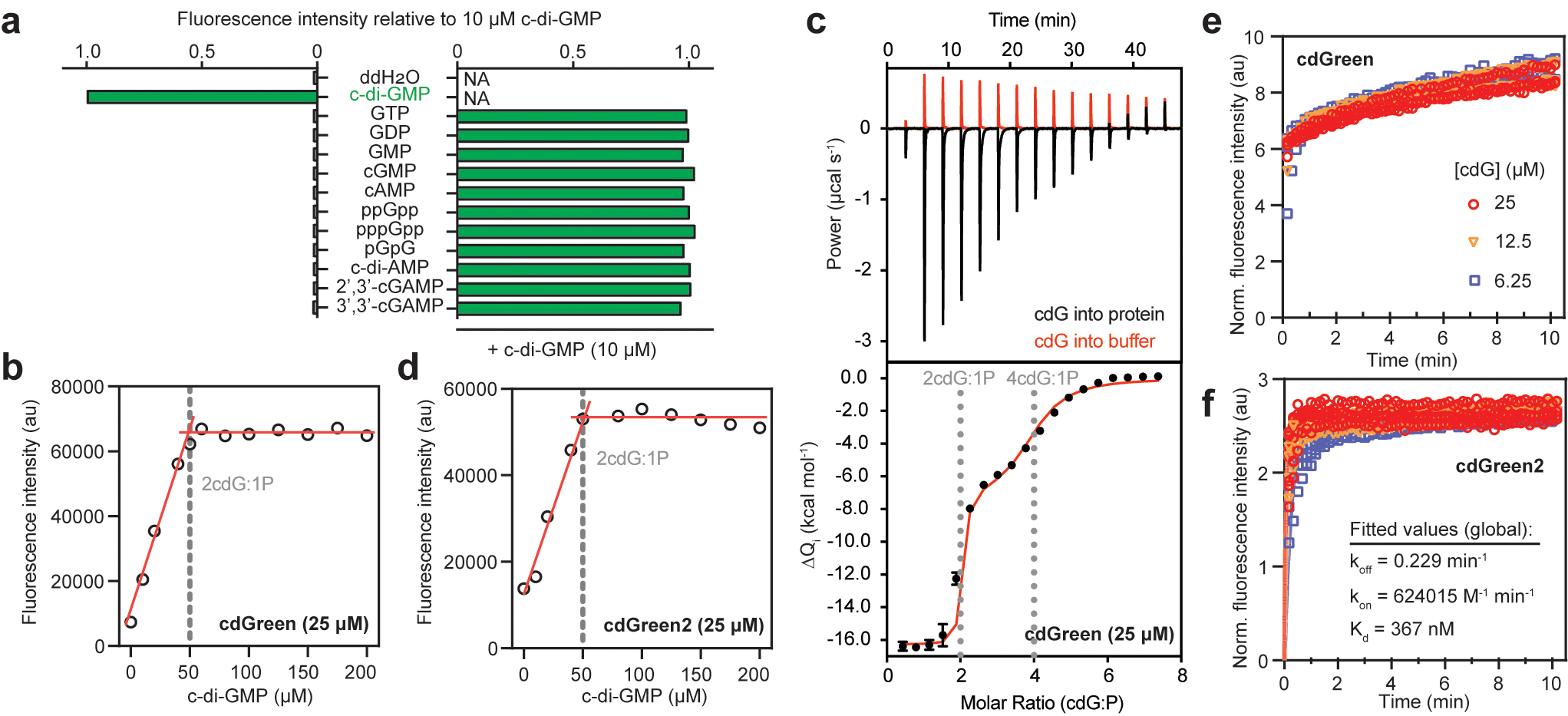

### Supplementary Figure 3

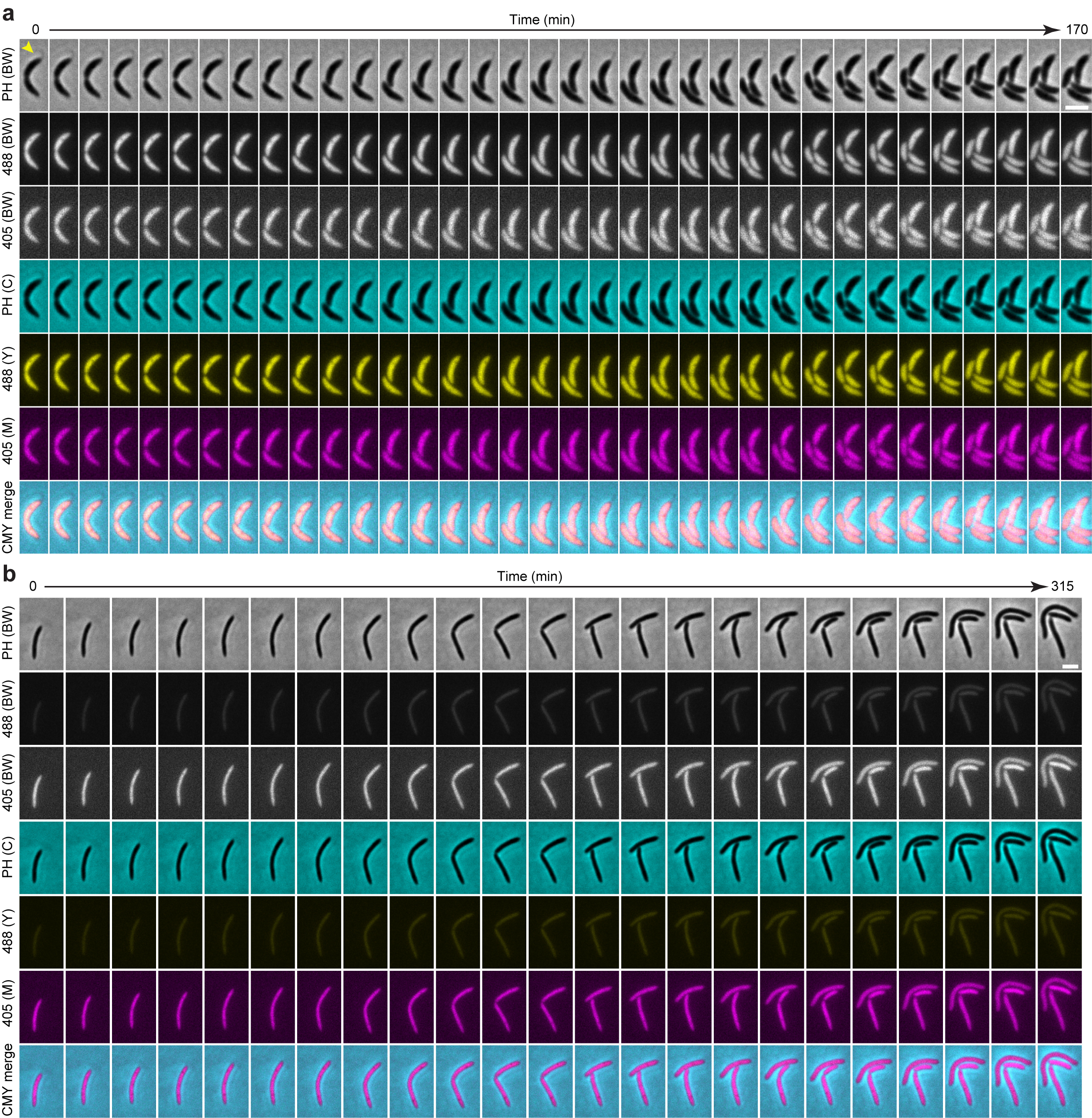

### Supplementary Figure 4

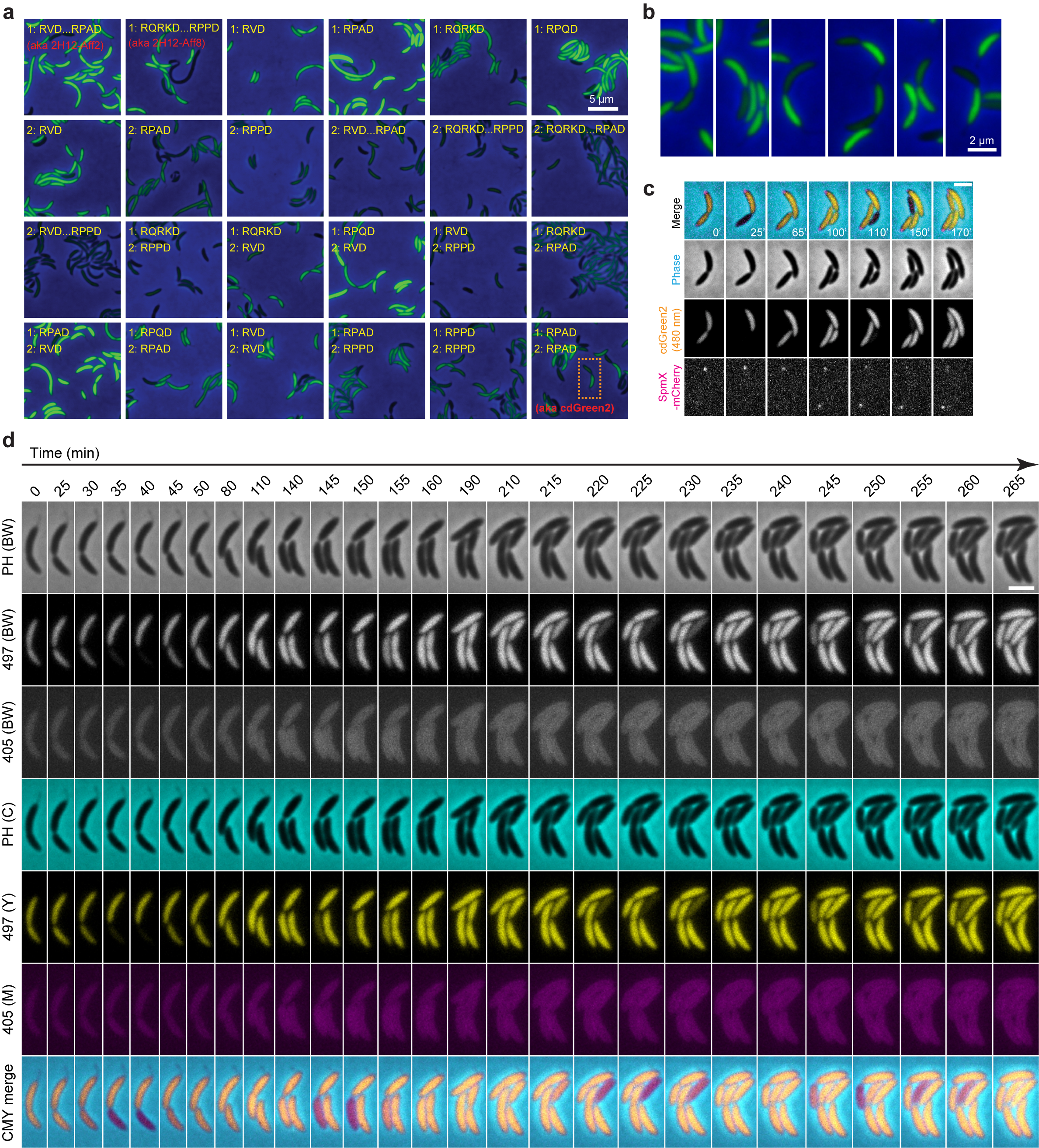

### Supplementary Figure 5

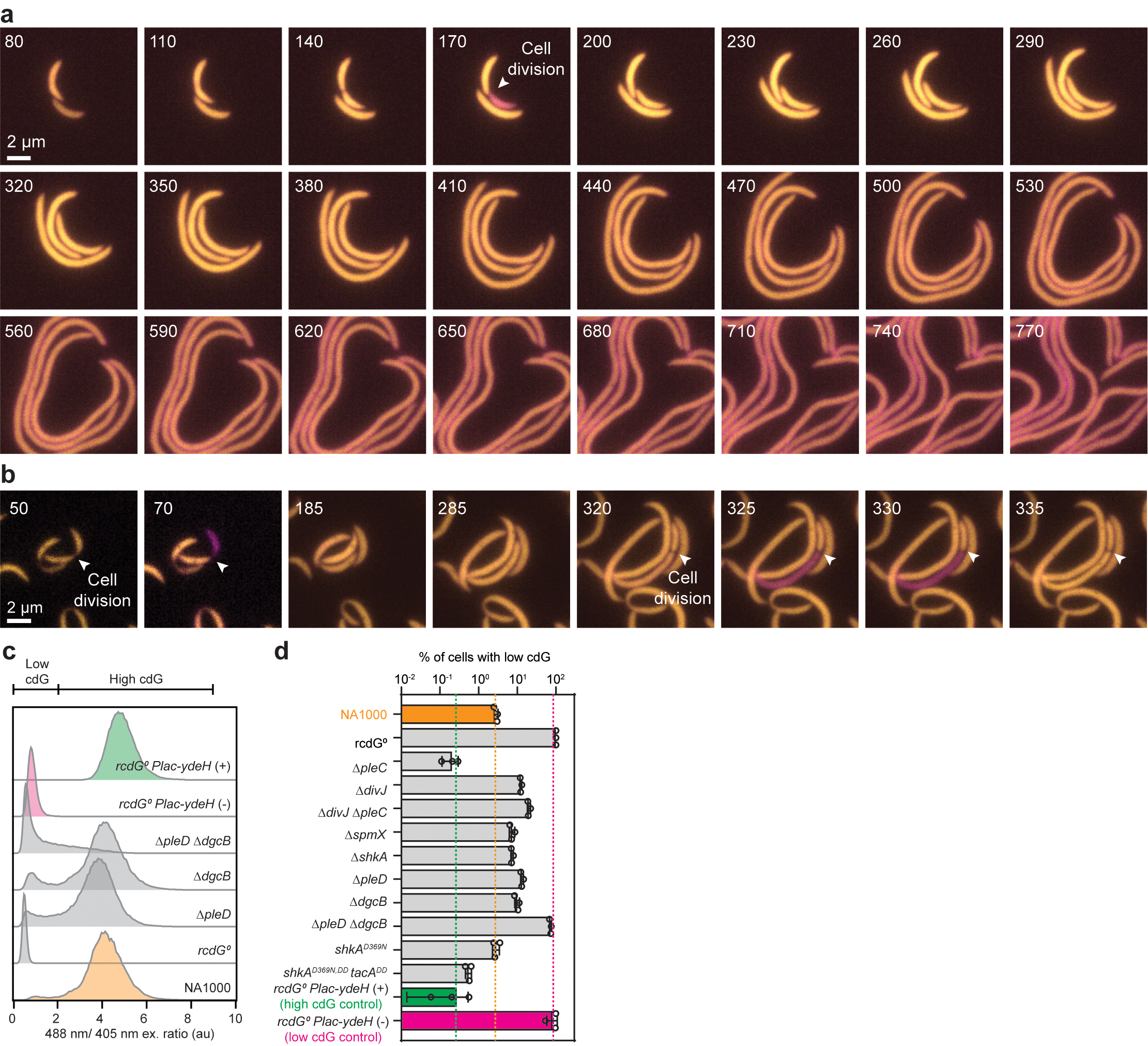

### Supplementary Figure 6

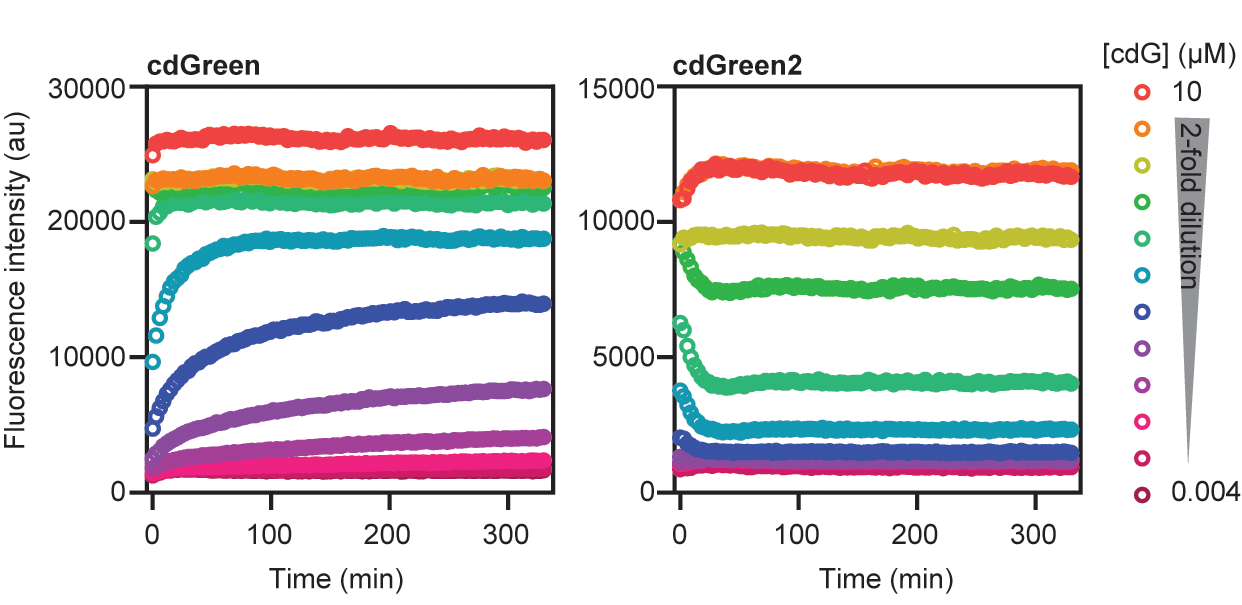
